# Optogenetic circuit mapping reveals connectivity and synaptic physiology of T-stellate projections from the cochlear nucleus to the auditory midbrain

**DOI:** 10.64898/2026.05.12.724651

**Authors:** Yoani N. Herrera, Britney M. Aguirre, Michael T. Roberts

## Abstract

T-stellate neurons in the ventral cochlear nucleus (VCN) receive synaptic input from the cochlear nerve and encode information about sound frequency and intensity, including rapid fluctuations in sound intensity that are important for speech processing. T-stellate neurons are the only neuron class in the VCN that projects directly to the inferior colliculus (IC), the midbrain hub of auditory processing. However, which IC neuron populations receive T-stellate input and how T-stellate input influences IC neuron excitability remain unknown. Using channelrhodopsin-assisted circuit mapping and whole-cell patch clamp recordings in brain slices, we compared the synaptic strength, prevalence, and short-term synaptic plasticity of T-stellate input to two molecularly defined classes of IC neurons: GABAergic neuropeptide Y (NPY) neurons and glutamatergic vasoactive intestinal peptide (VIP) neurons. Our results revealed that T-stellate neurons provide excitatory input to both NPY and VIP neurons, with T-stellate input to NPY neurons having a higher incident rate, larger magnitude, and faster kinetics than T-stellate input to VIP neurons. In many instances, T-stellate input also recruited feedforward inhibition and feedforward excitation onto NPY and VIP neurons. In addition, T-stellate input to NPY and VIP neurons exhibited short-term synaptic depression that became larger in amplitude at higher stimulation frequencies. These data provide insights on how T-stellate neurons influence individual neuron types and local circuits in the IC, laying a mechanistic foundation for investigating how T-stellate input contributes to frequency tuning, amplitude modulation selectivity, and speech processing in the IC.

## Introduction

Neural circuit function is regulated by the connection patterns between neuron types and the physiology of the synapses mediating those connections. In the central auditory system, most ascending projections from the lower auditory brainstem terminate in the midbrain at the inferior colliculus (IC), where a complex network of neurons integrate and process sound information before sending it to the thalamocortical system (Winer & Schreiner, 2005). However, due to the dense intermixing of axons in the IC (Oliver, 2005) and long-standing difficulty identifying IC neuron types (Drotos & Roberts, 2024), it is not clear which IC neuron types receive input from which auditory brainstem sources. It is also not clear how the physiology of synapses from specific ascending sources shapes neural computations in the IC.

T-stellate neurons in the ventral cochlear nucleus (VCN) are one of only three neuron types in the brain that both receive input from the cochlear nerve and project directly to the IC (Adams, 1979, 1983; Cant, 1982; Ryugo & Willard, 1985; Schofield & Cant, 1996; Oertel *et al*., 2011). Anatomical studies have shown that T-stellate neurons provide a large, tonotopically organized projection to the contralateral IC, with T-stellate axons ascending dorsomedially through the central nucleus of the IC (ICc) and reaching partway into the dorsal cortex of the IC (ICd) (Adams, 1979; Malmierca *et al*., 2005; Cant & Benson, 2008; Ryugo & Milinkeviciute, 2023). Physiologically, T-stellate neurons have relatively sharp tuning to sound frequency and convert the phasic firing of auditory nerve fibers into a sustained firing pattern that scales in firing rate as a function of sound intensity (Rhode & Smith, 1986; Young *et al*., 1988; Blackburn & Sachs, 1989). Because they are distributed across the tonotopic axis of the VCN, T-stellate neurons as a group provide a population code for the spectrum of sounds. Together, these properties make T-stellate neurons particularly well-suited for encoding fluctuations in the sound envelope, a critical feature for understanding speech (Oertel *et al*., 2011). However, surprisingly little is known about the connectivity and function of T-stellate synapses in the IC.

During repeated activation, many synapses exhibit short-term changes in synaptic strength, called short-term synaptic plasticity (STP) (Zucker & Regehr, 2002; Blitz *et al*., 2004; Regehr, 2012). Because STP shapes how information is passed between neurons, it plays a significant role in neural computations (Fortune & Rose, 2001; Abbott & Regehr, 2004; Chadderton *et al*., 2014; Larsen & Sjöström, 2015). However, how STP shapes computations in the IC has been challenging. First, axons from multiple ascending and descending sources overlap in the IC (Oliver *et al*., 1997; Coomes *et al*., 2005; Cant & Benson, 2006, 2008; Malmierca & Ryugo, 2011; Ryugo & Milinkeviciute, 2023), making it difficult to activate individual sources of input with electrical stimulation. Second, STP from a single presynaptic source can vary as a function of the postsynaptic neuron type (Larsen & Sjöström, 2015), but for many years, the field lacked reliable ways to target recordings to specific IC neuron classes (Drotos & Roberts, 2024). We are now well-positioned to overcome these challenges since, in recent years, our lab has identified vasoactive intestinal peptide (VIP) and neuropeptide Y (NPY) as markers for distinct types of glutamatergic and GABAergic neurons, respectively (Goyer *et al*., 2019; Silveira *et al*., 2020), and we showed that optogenetic circuit mapping can be used to selectively activate inputs from the cochlear nucleus to the IC (Goyer *et al*., 2019; Goyer & Roberts, 2020).

Here, we leveraged these advances to examine the connectivity and synaptic function of the T-stellate projection to the ICc. Based on the overlapping distributions of T-stellate axons, NPY neurons, and VIP neurons in the IC, we hypothesized that T-stellate neurons provide functional synaptic input to both these IC neuron classes. We further reasoned that T-stellate synapses onto NPY and VIP neurons would exhibit short-term depression (STD), as this would provide a potential mechanism to support the adaptation in firing rates IC neurons commonly exhibit in response to sustained sounds (Rose *et al*., 1963; Rees, 1992; Tan & Borst, 2007; Egorova *et al*., 2020). To test these hypotheses, we used optogenetic circuit mapping (Petreanu *et al*., 2007, 2009) to selectively activate T-stellate neuron terminals in the IC while making whole-cell recordings from NPY and VIP neurons in acutely prepared IC slices. Our results show that T-stellate neurons provide glutamatergic synaptic input to both NPY and VIP neurons, with input to NPY neurons being stronger and more common than that to VIP neurons. In many cases, activation of T-stellate projections also recruited local feedforward inhibitory and excitatory circuits in the IC. Additionally, we found that T-stellate synapses onto both neuron types exhibited similar levels of STD in response to trains of optical stimulation and that the extent of STD increased at higher stimulation frequencies. These results lay the foundation for future studies to identify how T-stellate input to specific IC circuits contributes to sound processing in vivo.

## Methods

### Animals

All experiments were approved by the University of Michigan Institutional Animal Care and Use Committee and were in accordance with NIH guidelines for the care and use of laboratory animals. Mice were kept on a 12-h day/night cycle with ad libitum access to food and water. To identify NPY neurons in the IC, Npy-IRES2-FlpO-D mice (B6.Cg-*NPY^tm1.1(flpo)Hze^*/J, Jackson Laboratory, stock #030211 (Daigle *et al*., 2018)) were crossed with Ai65F reporter mice (B6.Cg-*Gt(ROSA)26Sor^tm65.2(CAG-tdTomato)Hze^*/J, Jackson Laboratory, stock #032864 (Daigle *et al*., 2018)) to yield NPY-FlpO x Ai65F F1 offspring that expressed the fluorescent protein tdTomato in NPY neurons. To identify VIP neurons, VIP-IRES-Cre mice (B6J.Cg-*Vip^tm1(cre)Zjh^*/AreckJ, Jackson Laboratory, stock #031628 (Taniguchi *et al*., 2011)) were crossed with Ai14 reporter mice (B6.Cg-*Gt(ROSA)26Sor^tm14(CAG–tdTomato)Hze^*/J, Jackson Laboratory, stock #007914 (Madisen *et al*., 2010)) to yield VIP-Cre x Ai14 F1 offspring that expressed tdTomato in VIP neurons. All mice were on a C57BL/6J background, and because these mice present with early-onset age-related hearing loss (Noben-Trauth *et al*., 2003; Kane *et al*., 2012), mice used in experiments were aged postnatal day (P) 26-80 to limit potential effects of hearing loss. Mice of both sexes were used for all experiments.

### Intracranial Virus Injections

NPY-FlpO x Ai65F and VIP x Ai14 mice between the ages of P26-P48 were unilaterally injected with 40 nL of rAAV1-Syn-Chronos-GFP (Klapoetke *et al*., 2014) (Addgene #59170-AAV1, Lot #v123844, titer: 2.2 x 10^13^ GC/mL) or 80 nL of rAAV9-Syn-Chronos-GFP (University of North Carolina Vector Core, lot #av6102C, titer: 4.5×10^12^ GC/mL) into the right VCN. Virus injection surgeries were performed using standard aseptic techniques. Mice were induced with 4% vaporized isoflurane, then maintained throughout the procedure with 1-2% isoflurane, and their body temperature was maintained at 37°C using a homothermic heating pad. Mice were given an injection of the analgesic carprofen (5 mg/kg, Ostifen, MWI Animal Health, SKU# 510510), their scalps shaved, and an incision was made along the rostral-caudal axis of the scalp to expose the skull. Injection sites were mapped using stereotaxic coordinates relative to the skull surface at the lambda suture (485 µm caudal, 2455 µm lateral), and a craniotomy was drilled above the right VCN using a micromotor drill (K.1050, Foredom Electric Co.) equipped with a 0.5 mm burr (Fine Science Tools).

Viral constructs were injected into the brain with a NanoJect III nanoliter injector (Drummond Scientific Company) connected to an MP-285 micromanipulator (Sutter Instrument). Glass injection pipettes were prepared by pulling capillary glass (Drummond Scientific Company #3-000-203-G/X,) with a P-1000 microelectrode puller (Sutter Instrument). Glass injection pipettes were trimmed to 5 mm length and were back-filled with mineral oil before being front-filled with virus. Mice were injected at two deposit sites at 200 µm intervals along the dorsal-ventral axis of the VCN (4950-4750 μm deep from lambda), waiting 3-5 minutes between each deposit.

After injections were complete, the incision was closed using cyanoacrylate adhesive (3M VetBond) and treated with ∼0.5 mL of 2% lidocaine jelly. Mice recovered in a heated cage and returned to their home cage in the vivarium once completely ambulatory. Animals were checked daily for 7-10 days to track wound healing and were used for experiments ∼2-4 weeks following virus injection to allow sufficient time for Chronos expression.

### Brain Slice Preparation

Brain slices for patch-clamp electrophysiology were prepared from male and female mice aged P60–P80. Mice were deeply anesthetized with isoflurane, decapitated, and their brains were extracted quickly. Dissection of the IC was performed in 34°C artificial cerebrospinal fluid (ACSF) containing the following (in mM): 125 NaCl, 12.5 glucose, 25 NaHCO_3_, 3 KCl,1.25 NaH_2_PO_4_, 1.5 CaCl_2_,1 MgSO_4_, 3 sodium pyruvate, and 0.40 L-ascorbic acid, bubbled to a pH of 7.4 with 5% CO_2_ in 95% O_2_. Coronal IC sections (200 µm thick) were prepared using a vibrating microtome (VT1200S, Leica Biosystems). Slices were incubated at 34°C for 30 min in ACSF bubbled with 5% CO_2_ in 95% O_2_, then incubated at room temperature in the same ACSF for 30 min – 3 hours, before being transferred to the recording chamber. Dissections and slice incubation were performed in low light conditions, with the only illumination coming from red LEDs, to minimize unintended activation of optogenetic proteins.

### Analysis of Chronos-EGFP expression in the VCN and IC

To validate the location of injection sites and to assess viral spread, dissections for IC brain slice preparation were conducted to preserve the cochlear nucleus (CN). After IC slices were removed, the remainder of the tissue block containing the CN was drop-fixed overnight in 10% formalin and transferred to PBS the next day. Fixed tissue was stored at 4°C for up to 4 weeks, then coronal brain sections (50 µm thick) containing the CN were prepared using a vibrating microtome. VCN slices were mounted on slides, coverslipped with Fluoromount-G, and serial *z*-stack tile scan images of Chronos-EGFP expression throughout the CN were collected using a 20x objective on a Leica TCS SP8 laser scanning confocal microscope. The distribution of Chronos-EGFP positive neurons throughout the CN was assessed by comparison to the Allen Mouse Brain Reference Atlas (Allen Institute, n.d.) (RRID:SCR_002978).

### In-vitro Electrophysiological Recordings

For recordings, brain slices were transferred to a recording chamber under a fixed stage upright microscope (BX51WI, Olympus Life Sciences) where they were perfused at ∼2 ml/min with 34°C ACSF bubbled with 5% CO_2_ in 95% O_2_. The presence of T-stellate projections expressing Chronos-EGFP was confirmed in all IC slices using brief illumination of 470 nm light to visualize EGFP fluorescence. Recordings were targeted to NPY neurons identified in NPY-FlpO x Ai65F or VIP neurons identified in the VIP-Cre x Ai14 mice by the expression of tdTomato. Only neurons within the central nucleus of the IC (ICc) were selected for recording.

Recording electrodes were prepared from borosilicate glass capillaries (outer diameter 1.5 mm and inner diameter 0.86 mm; Sutter Instrument, #BF150-86-10) using a P-1000 microelectrode puller (Sutter Instrument). Electrode resistances ranged from 2.5 to 5.5 MΩ when filled with an internal solution consisting of the following (in mM): 115 K-gluconate, 7.73 KCl, 0.5 EGTA,10 HEPES,10 Na_2_-phosphocreatine, 4 MgATP, 0.3 NaGTP, and supplemented with 0.1% biocytin (wt/vol), with pH adjusted to 7.3 with KOH and osmolality to 290 mmol/kg with sucrose.

Current-clamp recordings were performed with a BVC-700A patch-clamp amplifier (Dagan Corporation). Current-clamp data were low-pass filtered at 10 kHz, sampled at 50 kHz with a National Instruments PCIe-6343 data acquisition board, and acquired using custom written algorithms in Igor Pro (Sutter Instrument). Voltage-clamp recordings were performed with a dPatch patch-clamp amplifier (Sutter Instrument). Voltage-clamp data were low-pass filtered at 10 kHz, sampled at 50 kHz, and acquired using SutterPatch software (Sutter Instrument). Bridge balance was corrected in all current-clamp recordings, and series resistance and pipette capacitance compensation were applied in all voltage-clamp recordings using 80-98% correction. All membrane potentials have been corrected post hoc for a liquid junction potential of 11 mV. Recordings with series resistance >25 MΩ were discarded.

Input resistance was assessed by finding the slope of the voltage vs current relationship measured at the peak (R_pk_) and steady-state (R_ss_) portion of responses to a series of 100 ms current steps delivered in 10 pA increments that hyperpolarized the membrane potential between −15 and 0 mV relative to rest. To determine membrane time constant, we applied 50 current steps that hyperpolarized the membrane potential by 2–6 mV, fit a single exponential function to each response, and then calculated the median time constant from the exponential fits.

A minimum stimulation paradigm was used to elicit optogenetic activation of T-stellate projections in the IC. For this, short bursts of blue light were delivered through the 40x microscope objective using a 470 nm LED coupled to the microscope’s epifluorescence path. The LED was adjusted to the lowest intensity that elicited an EPSC or EPSP in most but not all light presentations. The resulting optical power densities ranged from 4-56 mW/mm^2^, as measured by a S130C photodiode power sensor (ThorLabs) connected to a PM100D power meter (ThorLabs). Light pulses ranged from 0.25 ms – 10 ms but were generally 1-5 ms. For off-bouton excitation experiments, the 40x objective was moved ∼430 µm away from the center of the recorded soma, such that the field of illumination was completely off the recorded neuron.

To isolate EPSCs, the ACSF for all voltage-clamp experiments contained 5 µM SR95531 (gabazine, GABA_A_ receptor antagonist, HelloBio # HB0901) and 1 µM strychnine hydrochloride (glycine receptor antagonist, Millipore-Sigma #S8753). To determine the receptors mediating T-stellate input to IC neurons, NMDA receptors were blocked using 50 µM D-AP5 (HelloBio # HB0225) and AMPA receptors were blocked using 10 µM NBQX disodium salt (HelloBio # HB0443). In some experiments, we tested for monosynaptic input using 0.1 mM tetrodotoxin citrate (TTX, sodium channel blocker, HelloBio, #HB1035) and 1 mM 4-AP (K^+^ channel blocker, Sigma-Aldrich, #275875) dissolved in high Ca^2+^ (2.5 mM) ASCF (Petreanu *et al*., 2009).

Analyses of electrophysiology data were performed using custom algorithms in Igor Pro. Postsynaptic response amplitude, half-width, latency, and jitter were measured, then averaged for each cell. Data from voltage-clamp experiments were digitally low-pass filtered at 5 kHz prior to analysis.

### Post hoc reconstruction of neuron morphology and morphology analysis

During whole-cell recordings, neurons were filled with biocytin (Thermo Fisher Scientific, catalog # B1592) via the recording pipette. After the recording, the pipette was slowly removed to allow the cell membrane to reseal, and the brain slice was fixed overnight with 10% formalin and transferred to PBS the next day. Slices were stored at 4°C for up to 4 weeks, then stained using biocytin-streptavidin histochemistry. Slices were washed in PBS 3 x 10 minutes each, tissue was permeabilized using 0.2% Triton X-100 in PBS for 2 h, washed again 3 x 10 min in PBS, and then stained for 24 h with streptavidin-Alexa Fluor-647 (1:1000, Thermo Fisher Scientific, catalog # S21374) at 4°C. The following day, slices were again washed 3 x 10 min with PBS, fixed in 10% formalin for 1 h, and then washed 3 x 10 min with PBS. Slices were then mounted on slides and cover-slipped using Fluoromount-G. *z*-stack tile scan images of Chronos-EGFP-labelled T-stellate projections and streptavidin-Alexa Fluor-647 stained neurons in the IC were collected using a 1.4 NA 63x oil-immersion objective on a Leica TCS SP8 laser scanning confocal microscope. To assess whether T-stellate synapses were adjacent to recorded NPY and VIP neuron dendritic arbors, we imaged regions of interest where EGFP and Alexa Fluor-647 were in close proximity.

### Statistics

Statistical analyses were performed using RStudio (RStudio 2024.12.1+563, Boston) for R 4.4.2 (R Core Team, 2026), and effects were considered significant when p < 0.05. To compare the incidence of T-stellate input to NPY and VIP neurons, we used a two sample proportions test (R: prop.test) and report χ^2^ and p values. To compare two groups for the intrinsic physiology parameters, we used a two-sample Welch’s t-test (R: t.test) and report t-statistics and p values. To compare multiple groups for the receptor pharmacology and stimulus train experiments, we used linear mixed model (LMM) analyses (R: ‘lme4,’ ‘lmerTest,’ and ‘emmeans’ packages (Bates *et al*., 2015; Kuznetsova *et al*., 2017; Lenth & Piaskowski, 2025)), and report β values, 95% confidence intervals, and p values. Lowercase “n” indicates the number of neurons recorded for an experiment, and capital “N” indicates the number of animals from which those neurons were recorded. Summary statistics are shown as mean ± SD, unless otherwise indicated.

## Results

### Expressing the excitatory channelrhodopsin, Chronos, in T-stellate projections to the IC

Because the axons of ascending auditory projections overlap extensively in the IC, it is not experimentally tractable to use electrical stimulation to activate axons from a single, identified input source in IC brain slices. Optogenetic circuit mapping provides a way around this problem so long as the channelrhodopsin can be selectively expressed in neurons from one input source. We therefore tested whether we could selectively express the fast, excitatory channelrhodopsin Chronos (Klapoetke *et al*., 2014) in the axons and terminals of T-stellate projections to the IC. As T-stellate neurons are the only VCN neurons that project to the IC, we used stereotaxic surgeries to inject an AAV encoding Chronos-EGFP with a pan-neuronal promoter (AAV1-hSyn-Chronos-EGFP) into the VCN of transgenic mice that express tdTomato in NPY or VIP neurons (Npy-IRES2-FlpO-D x Ai65F mice and VIP-IRES-Cre x Ai14 mice, respectively; **Figure 1A**).

**Figure 1.**
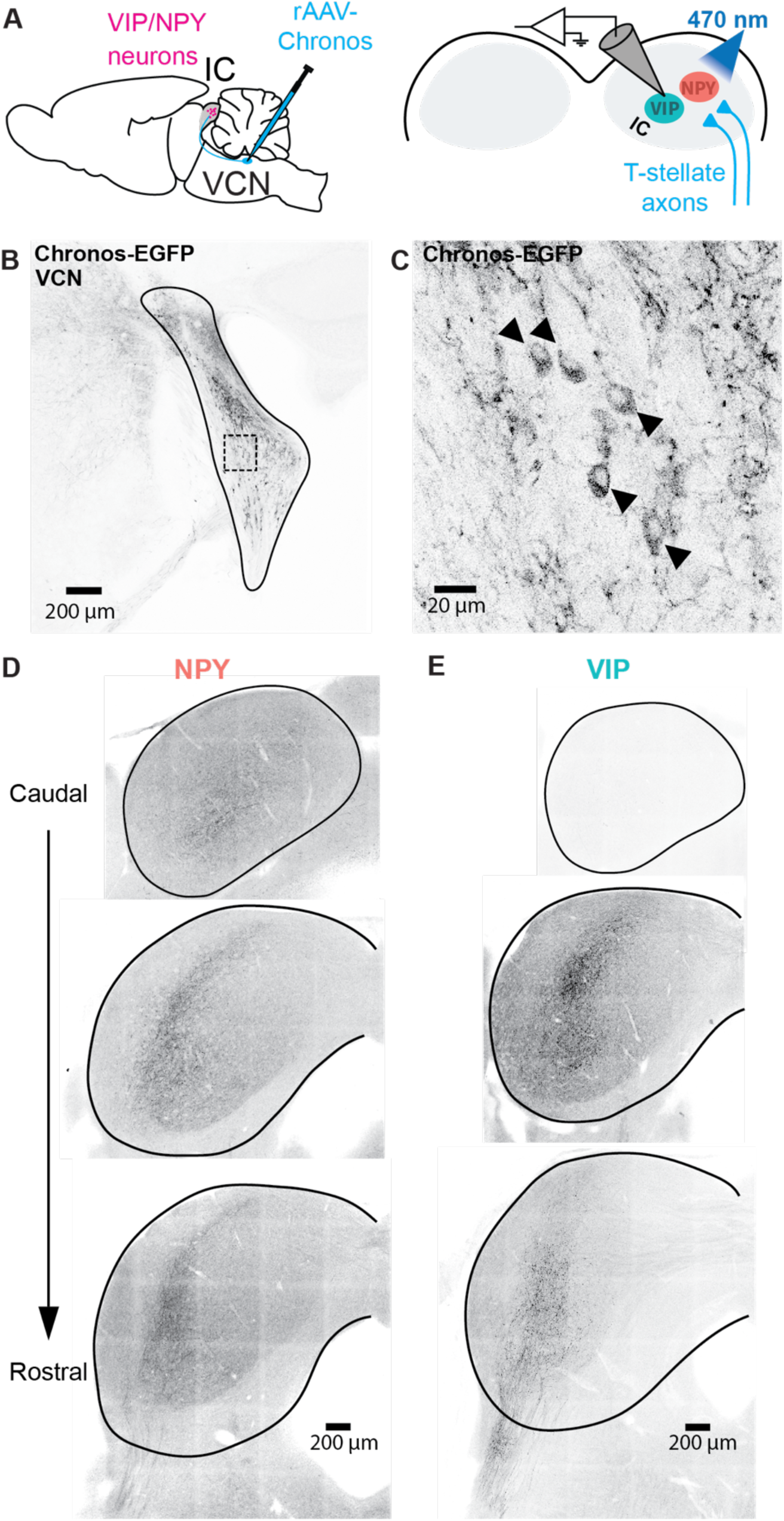
Expression of Chronos-EGFP in T-stellate projections to the contralateral IC. **A.** Schematic of experimental setup. An AAV1 encoding Chronos-EGFP under control of the synapsin promoter was injected into the right VCN. Recordings were targeted to tdTomato+ NPY or VIP neurons in brains slices prepared from the IC contralateral to the injection site. **B.** Example image showing Chronos-EGFP expression in the VCN. AAV injection sites were validated in 17 NPY and 16 VIP mice. In all cases, Chronos-EGFP+ somata were observed in the VCN. **C.** Higher magnification view of the VCN region outlined by a dashed line in **B**. Chronos-EGFP+ somata are highlighted with arrowheads. **D,E.** 4 NPY and 4 VIP mice were perfused to examine the distribution of Chronos-EGFP+ axons and terminals across the IC. The distribution of Chronos-EGFP+ in both NPY (**D**) and VIP (**E**) mice was consistent with the distribution of T-stellate projections reported in previous studies (Adams, 1979; Malmierca *et al*., 2005; Cant & Benson, 2008; Ryugo & Milinkeviciute, 2023). Images show example IC coronal sections arranged from caudal (top) to rostral (bottom). In **B-E**, epifluorescence images are presented in inverted grayscale.

We observed Chronos-EGFP labeled axons and terminals forming a band that extended up through the central nucleus of the contralateral IC and entered the ventral edge of the dorsal cortex of the contralateral IC (ICd), consistent with T-stellate projection patterns described in previous tract tracing studies (Adams, 1979; Malmierca *et al*., 2005; Cant & Benson, 2008; Ryugo & Milinkeviciute, 2023) (**Figure 1B**). Injection sites were examined in 21 Npy-IRES2-FlpO-D x Ai65F mice and 18 VIP-IRES-Cre x Ai14 mice, hereafter referred to as NPY and VIP mice, respectively. Chronos-EGFP expressing cells were found in all VCN slices imaged (**Figure 1C,D**). Viral transduction appeared to work equally well in NPY and VIP mice, and in both, the distribution of T-stellate axons varied across the rostral-caudal axis of the IC, again consistent with previous anatomical studies (Adams, 1979; Malmierca *et al*., 2005; Cant & Benson, 2008; Ryugo & Milinkeviciute, 2023) (**Figure 1E,F**).

Spread of virus beyond the VCN into the dorsal cochlear nucleus (DCN) could confound results, since fusiform and giant cells in the DCN also project to the IC (Ryugo & Willard, 1985). To minimize virus spread into the DCN, our stereotaxic injection coordinates were optimized to target the rostral VCN, away from the DCN, which is situated more caudally. Supporting the efficacy of this approach, the pattern of Chronos-EGFP labeled axons observed in the contralateral IC was consistent with VCN projections, but not DCN projections, which branch into the IC lateral cortex (IClc) at angles oblique to the main projection into the ICc (Ryugo & Milinkeviciute, 2023) (**Figure 1B,E, F**).

In addition to examining VCN injection sites, we examined Chronos-EGFP expression patterns throughout the entire cochlear nucleus (CN) in 9 NPY and 8 VIP mice. In all CN image series, the most rostral slices contained Chronos-EGFP positive neurons, which were likely a mix of VCN neuron types, including T-stellate cells (**Figure 2**, leftmost panels). Sections from the mid-rostral-caudal CN contain both VCN and DCN. We routinely saw transduced cells in the posterior VCN in these sections, which were likely a mix of T-stellate cells, D-stellate cells, and octopus cells, as well as some transduced cells in the deep layer of the ventral DCN. Sections containing the DCN also showed Chronos-EGFP-labeled axons and terminals, consistent with T-stellate and/or D-stellate cell projections from the VCN into the DCN, but across all sections examined from 17 mice, only two labeled cell bodies were observed in the fusiform cell layer or molecular layer of the DCN (**Figure 2**, rightmost panels). These results suggest that AAV injections were successfully targeted to the VCN and that the virally labeled somata occasionally observed in deep layers of the DCN were most likely vertical cells (tuberculoventral cells) retrogradely labeled through their axonal projections to the VCN (Wickesberg & Oertel, 1988; Wickesberg *et al*., 1991). While we cannot completely rule out the possibility that a small number of DCN neurons that project to the IC (fusiform and/or giant cells) were transduced by virus injections in some mice, our anatomical data suggest that off-target labeling of IC projection neurons was rare and unlikely to significantly impact optogenetic circuit mapping experiments. Thus, since T-stellate neurons are the only VCN neurons that project to the IC, stereotaxic injection of a pan-neuronal AAV in the VCN provides an effective way to target Chronos-EGFP expression to T-stellate axons for optogenetic circuit mapping experiments in the IC.

**Figure 2.**
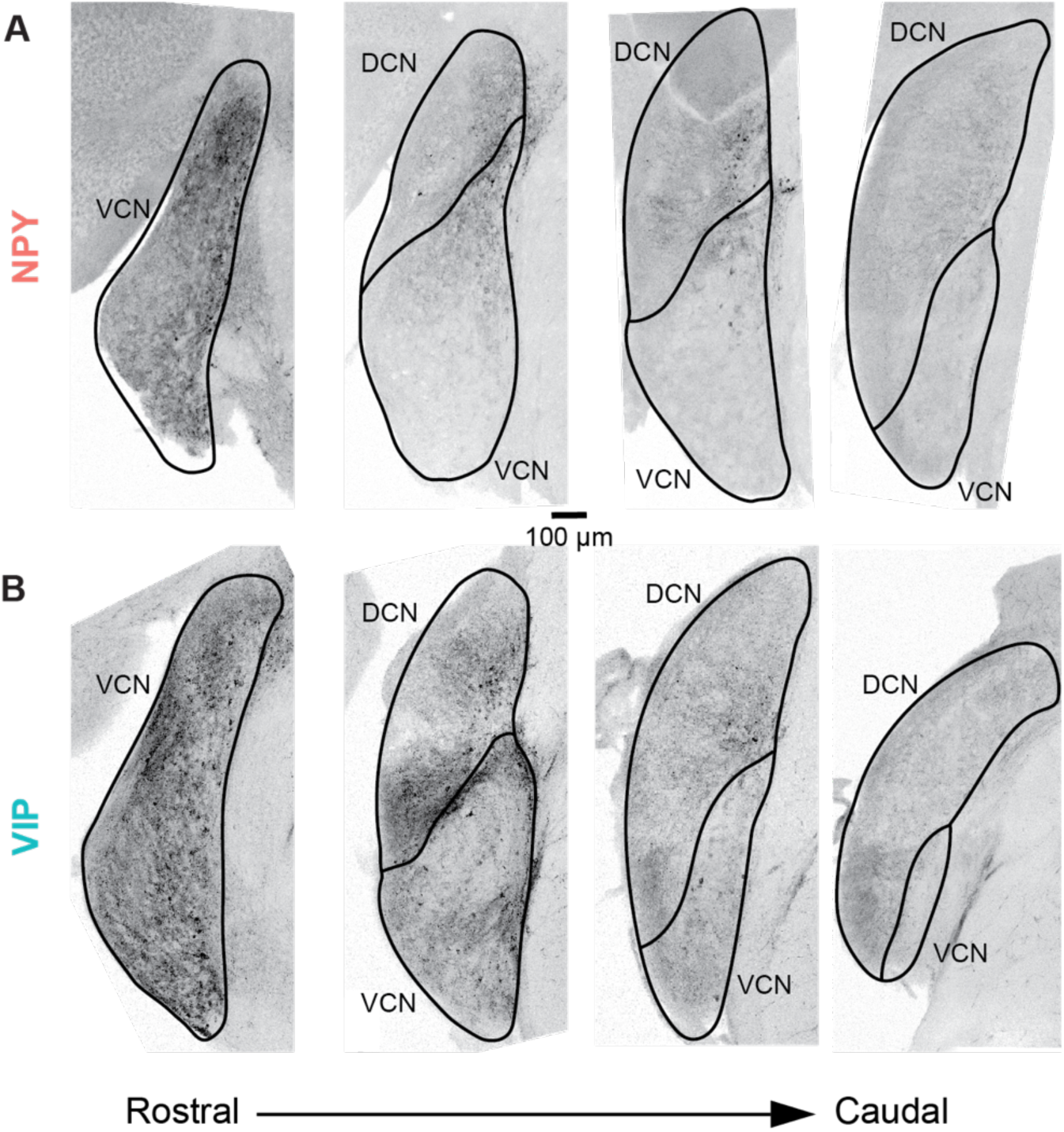
Stereotaxic AAV injections led to strong expression of Chronos-EGFP in VCN neurons with little to no expression in the DCN. **A,B.** In 9 NPY and 8 VIP mice, we examined the distribution of Chronos-EGFP+ cells across the entire CN. Epifluorescence images show a series of coronal sections arranged from rostral (left) to caudal (right) from the cochlear nucleus of an NPY mouse (**A**) and a VIP mouse (**B**). The VCN contained numerous Chronos-EGFP+ somata, while very few Chronos-EGFP+ somata were observed in the DCN. When labeled somata were observed in the DCN, they were almost exclusively located in the DCN deep layer and not in the fusiform cell layer, where most DCN principal neurons are located. Sparse labeling in the deep layer of the DCN is consistent with a retrograde labeling of small numbers of DCN vertical cells via their projections to the VCN. Chronos-EGFP+ axons were also routinely observed in the DCN, many of which likely originated from T-stellate and D-stellate neurons in the VCN. Epifluorescence images are presented in inverted grayscale.

### T-stellate neurons provide synaptic input to NPY and VIP neurons in the ICc

To test whether NPY and VIP neurons receive functional input from T-stellate neurons in the VCN, we used channelrhodopsin-assisted circuit mapping to selectively activate T-stellate synapses in the ICc while targeting whole-cell recordings to NPY or VIP neurons in acutely prepared IC brain slices. For all recordings, tdTomato+ NPY or VIP neurons located in the ICc were targeted based on their proximity to EGFP+ T-stellate terminals. To activate T-stellate terminals, we used a minimum stimulation paradigm in which the LED power was set to elicit postsynaptic responses in most, but not all, presentations of light. Across all experiments, we recorded from total of 178 NPY neurons and 242 VIP neurons. A significantly larger portion of NPY neurons received functional input from T-stellate neurons; excitatory postsynaptic responses were observed in 53% of NPY neurons and 29% of VIP neurons (n = 94/178 NPY and 70/242 VIP, χ^2^ =25.58, p < 0.001, two sample proportions test).

The NPY and VIP neurons recorded from during current-clamp recordings had intrinsic physiological properties similar to those reported in previous studies (Goyer *et al*., 2019; Silveira *et al*., 2020, 2024). We found that there was little to no difference in resting membrane potential, peak input resistance (R_PK_), steady state input resistance (R_ss_), or membrane time constant (τ) between the neurons in this experiment and those in previous reports from our lab (**Table 1**).

**Table 1.** The physiological properties of NPY and VIP neurons were similar to those reported in previously published studies. Comparisons were made to results from: (Goyer *et al*., 2019; Silveira *et al*., 2020).

|  | Recorded NPY | Published NPY | t-value | df | p-value | Recorded VIP | Published VIP | t-value | df | p-value |
| --- | --- | --- | --- | --- | --- | --- | --- | --- | --- | --- |
| Resting Membrane Potential (mV) | $-66.8 \pm 3.9$ | $-65.6 \pm 5.1$ | -1.17 | 69.8 | 0.24 | $-67.6 \pm 3.4$ | $-70.5 \pm 4.4$ | 3.59 | 32.2 | 0.001 |
| Steady state input resistance ($R_{ss}$, MΩ) | $229 \pm 126$ | $219 \pm 103$ | 0.36 | 58.7 | 0.72 | $331 \pm 190$ | $242 \pm 139$ | 2.18 | 25.7 | 0.04 |
| Peak input resistance ($R_{peak}$, MΩ) | $196 \pm 127$ | $212 \pm 130$ | -0.56 | 47.9 | 0.58 | $301 \pm 189$ | $237 \pm 171$ | 1.51 | 24.3 | 0.14 |
| Membrane time constant ($\tau$, ms) | $10.6 \pm 6.5$ | $13.2 \pm 7.4$ | -1.73 | 64.3 | 0.09 | $19.7 \pm 8.5$ | $15.0 \pm 8.8$ | 2.5 | 26.8 | 0.02 |

We next investigated the basic properties of optically evoked excitatory postsynaptic potentials (oEPSPs) between NPY and VIP neurons. In current-clamp recordings, oEPSPs in NPY neurons (n = 30, N = 20) had mean amplitudes of 3.28 mV ± 1.72 mV and mean half-widths of 8.43 ± 4.21 ms, while those in VIP neurons (n = 24, N = 12) had mean amplitudes of 1.62 ± 1.23 mV and mean half-widths of 15.94 ± 12.22 ms (**Figure 3A**). oEPSPs in NPY neurons exhibited significantly larger amplitudes and smaller half-widths that those in VIP neurons (amplitude: t = −11.45, df = 543.17, *p* <0.001; half-width: t = −2.87, df = 27.39, *p* = 0.007, Welch’s t-test) (**Figure 3B**). Since T-stellate neurons are glutamatergic (Cao *et al*., 2019), we used pharmacology to test whether oEPSPs in NPY and VIP neurons were mediated by AMPA and/or NMDA receptors. We found that the NMDA receptor antagonist D-AP5 (50 µM) slightly increased EPSP amplitudes in NPY neurons (control = 3.35 ± 0.21 mV, D-AP5 = 4.10 ± 0.28 mV, control vs AP5: β = 0.75, 95% CI (−1.36, −3.18), *p* = 0.02, linear mixed model (LMM); n = 9, N = 9), but did not alter EPSP amplitudes in VIP neurons (control = 1.91 ± 0.23 mV, D-AP5 = 1.94 ± 0.30 mV, LMM for control vs AP5: β = 0.03, 95% CI (−0.64, 0.58), p = 0.99; n = 8, N = 6). The AMPA receptor antagonist NBQX (10 µM) blocked oEPSPs in both NPY and VIP neurons (NPY: NBQX = 0 ± 0.0 mV, LMM for control vs NBQX: β = 3.35, 95% CI (2.74, 3.97), *p* < 0.001; VIP: NBQX = 0 ± 0.0 mV, LMM for control vs NBQX: β = 1.91, 95% CI (1.30, 2.52), *p* < 0.001) (**Figure 3C-D**). Together, these data indicate that T-stellate neurons provide excitatory synaptic input to both NPY and VIP neurons in the ICc, with postsynaptic responses primarily mediated through AMPA receptors. T-stellate input to NPY neurons, however, had a higher incident rate, larger EPSP amplitude, and faster EPSP kinetics than T-stellate input to VIP neurons.

**Figure 3.**
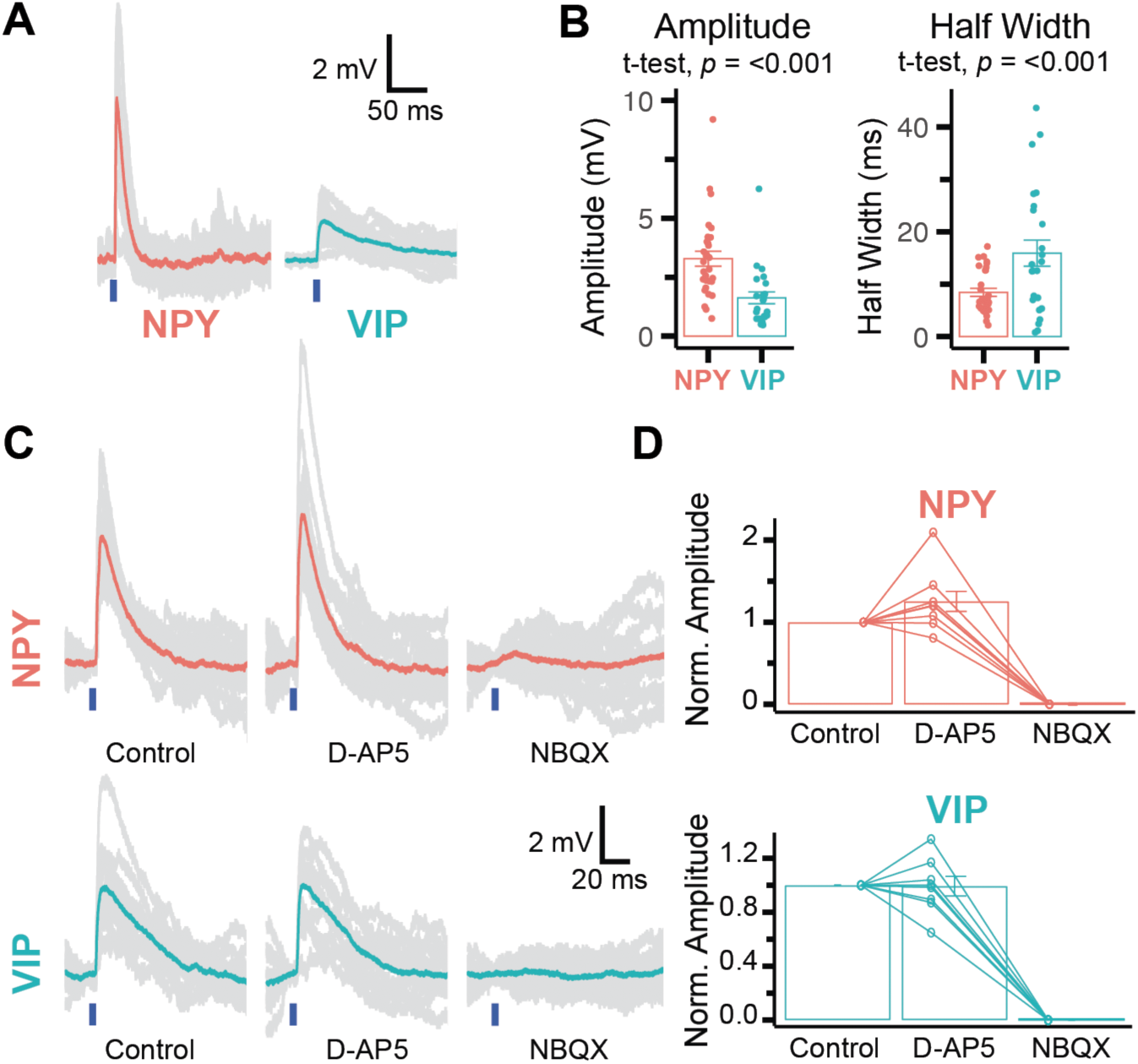
T-stellate neurons provide excitatory synaptic input to NPY and VIP neurons. **A.** Examples of EPSPs elicited in an NPY and a VIP neuron through optogenetic stimulation of T-stellate projections (oEPSPs). Blue bars indicate when optical stimuli (light pulses) were presented. **B.** oEPSPs elicited in NPY neurons had greater amplitudes and faster kinetics than those elicited in VIP neurons (n = 30 NPY, 24 VIP). **C,D.** In both NPY neurons (top row) and VIP neurons (bottom row), oEPSPs were not significantly altered by application of the NMDA receptor antagonist D-AP5 (50 µM), but were completely blocked by the AMPA receptor antagonist NBQX (10 µM).

### Anatomical support for T-stellate neuron input to NPY and VIP neurons

We next examined whether NPY and VIP neurons that exhibited oEPSPs were located adjacent to T-stellate neuron terminals. During whole-cell recordings, neurons were dialyzed through the recording pipette with an internal solution containing 0.1% biocytin. This allowed for post hoc reconstruction of the morphology of recorded neurons and mapping of Chronos-EGFP+ T-stellate terminals located adjacent to the dendrites and somata of those neurons. We validated that streptavidin-Alexa Fluor 647-labeled neurons were tdTomato+ (**Figure 4A**, left panel inset) and then found that labeled neurons were located in regions that had many Chronos-EGFP+ axons and terminals (**Figure 4A,C**). Under higher magnification (1.40 NA 63x objective), Chronos-EGFP+ terminals were routinely observed close in proximity to the dendritic arbors of both NPY and VIP neurons that exhibited oEPSPs (**Figure 4B,D**). Thus, these anatomical results support the conclusion that the NPY and VIP neurons that exhibited oEPSPs received synaptic input from T-stellate neurons.

**Figure 4.**
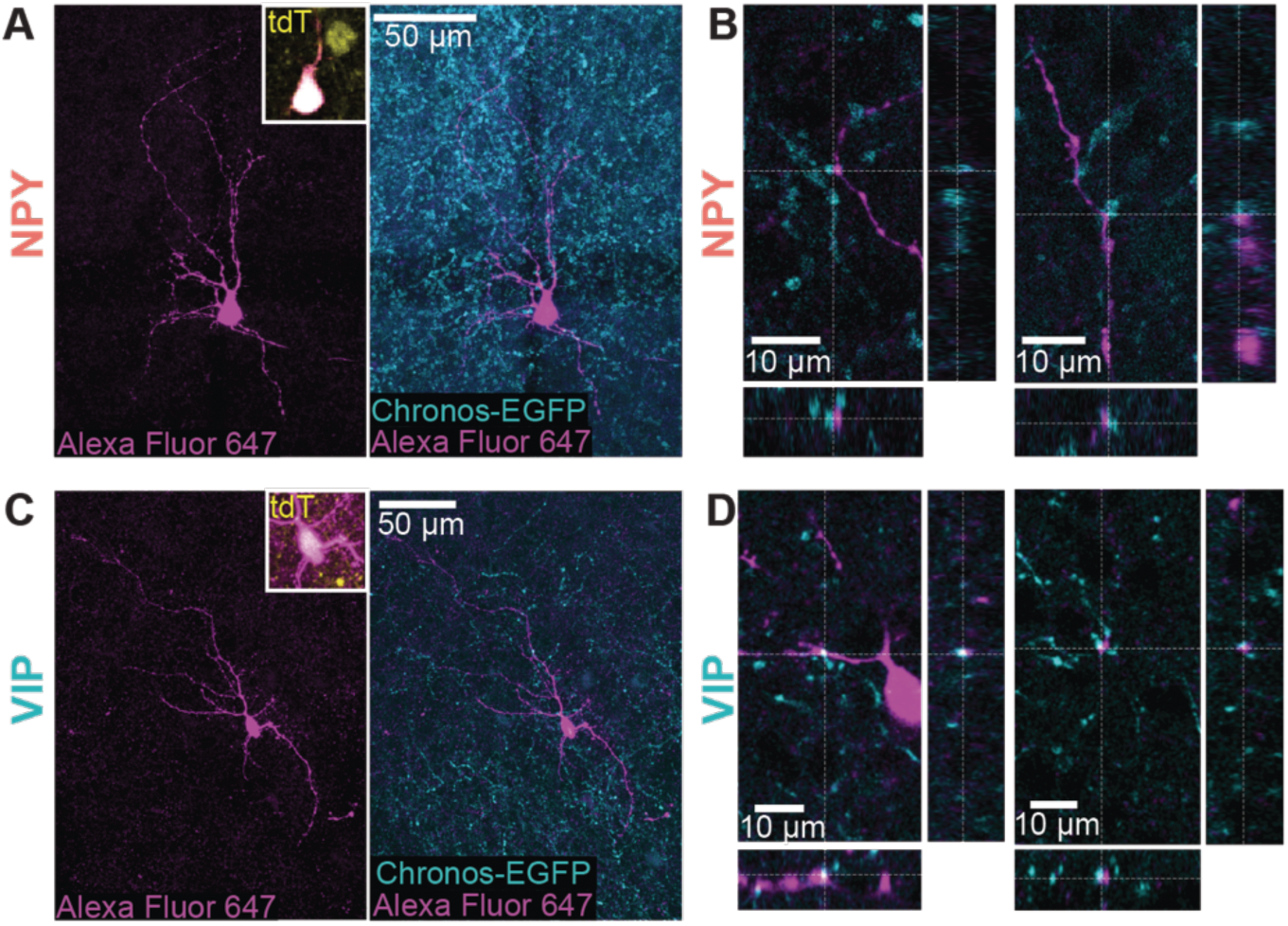
Post hoc imaging showed Chronos-EGFP+ puncta close in proximity to the dendrites of recorded NPY and VIP neurons. During recordings, neurons were dialyzed with an internal solution containing biocytin. At the end of recordings, slices were fixed, and recorded neurons were stained with streptavidin-Alexa Fluor 647. **A.** (left) Example showing the morphology of a streptavidin-Alexa Fluor 647-labeled NPY neuron that exhibited oEPSPs during the recording. Inset shows overlay with tdTomato, confirming that the neuron was a tdTomato+ NPY neuron. (right) Chronos-EGFP+ puncta were abundant in the region surrounding the recorded NPY neuron. **B.** Higher magnification images showing Chronos-EGFP+ puncta close in proximity to two example dendritic regions from the NPY neuron shown in **A**. Each dendritic region is shown from three vantage points. The large image shows a maximum intensity projection along the z-axis (depth-axis), and the smaller images show maximum intensity projections along the x-axis and y-axis. The intersection of dashed lines indicates where a Chronos-EGFP+ puncta colocalized with the dendrite of the recorded neuron in all three image planes. **C,D.** Example of a streptavidin-Alexa Fluor 647-labeled VIP neuron with dendrites located close in proximity to Chronos-EGFP+ puncta. Images are laid out as in **A,B**.

### T-stellate neurons can recruit feedforward circuits in the IC

In many of the recordings performed for this study, individual light pulses elicited a single oEPSP or optically evoked excitatory postsynaptic current (oEPSC) in NPY and VIP neurons. These single events had low latency and jitter, consistent with direct activation of T-stellate projections synapsing on the recorded cell (i.e., monosynaptic connections; **Table 2**). To confirm that short latency events were monosynaptic, we used voltage-clamp recordings to compare oEPSCs under control conditions and after applying 0.5 µM tetrodotoxin (TTX, Na^+^ channel blocker) and 1 mM 4-aminopyridine (4-AP, K^+^ channel blocker) to isolate monosynaptic inputs (Petreanu *et al*., 2009). We also increased the Ca^2+^ concentration in the ACSF to 2.5 mM throughout this experiment to enhance release probability. We found that blocking polysynaptic pathways did not abolish oEPSCs in most NPY and VIP neurons (n = 7/9 NPY; n = 8/9 VIP) (**Figure 5A**).

**Figure 5.**
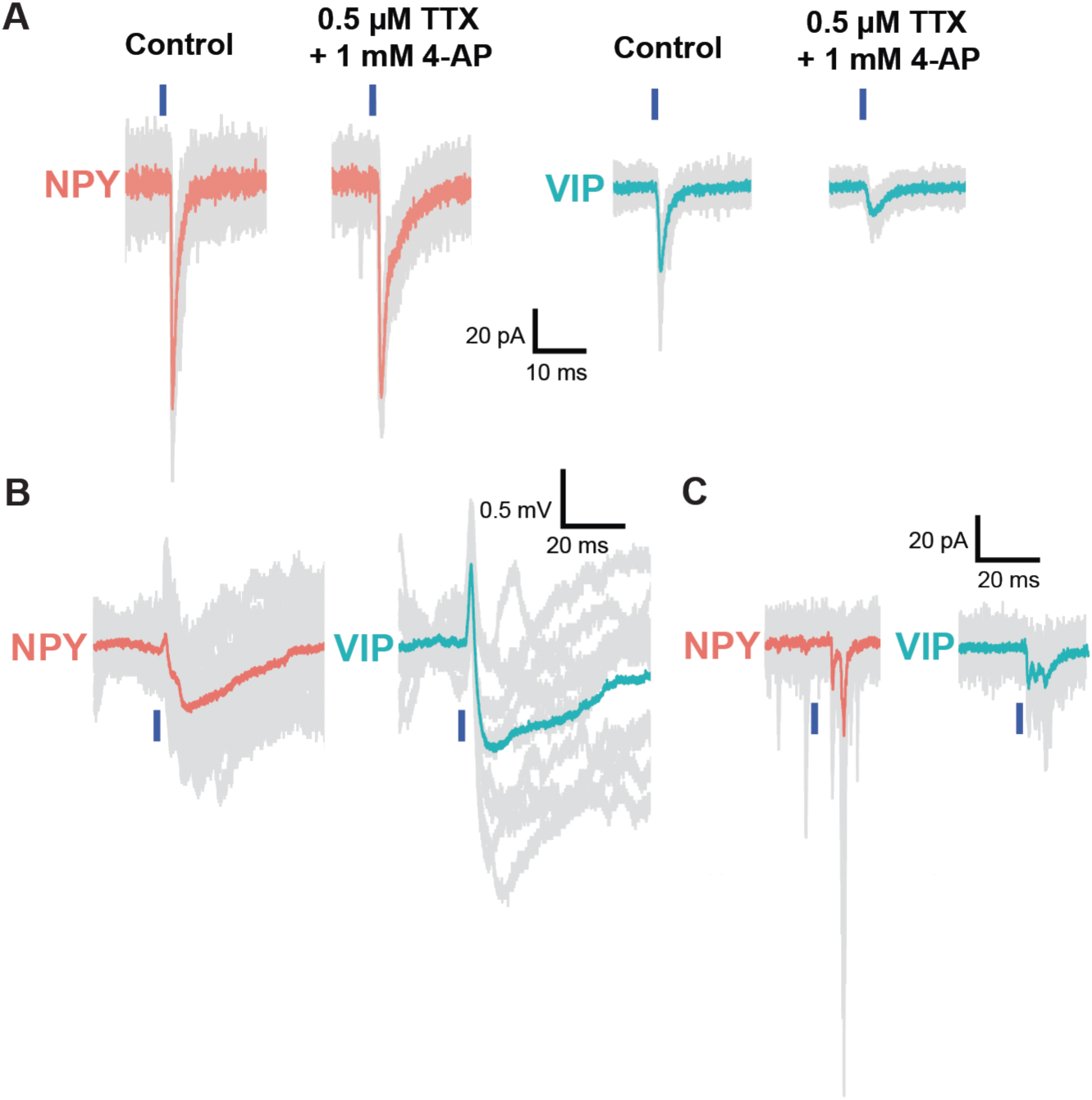
Optogenetic stimulation of T-stellate projections can recruit local feedforward circuits in the IC. **A.** In voltage-clamp recordings, short-latency oEPSCs in NPY and VIP neurons were resistant to block with 0.5 µM TTX, indicating that these oEPSCs were elicited by direct input from T-stellate synapses, not feedforward projections. To enhance synaptic release for the TTX experiment, the ACSF Ca^2+^ concentration was raised to 2.5 mM throughout the TTX experiment, and TTX was bath-applied in combination with 1 mM 4-AP. **B.** In both NPY and VIP neurons (n = 6 NPY and n = 6 VIP), stimulation of T-stellate afferents could elicit an initial, direct oEPSP followed by longer-latency IPSPs. Since the brain slice contained only IC neurons, the longer-latency events were likely due to T-stellate activation of local, feedforward circuits. **C.** Similarly, stimulation of T-stellate afferents elicited longer-latency EPSCs in several NPY and VIP neurons (n = 19 NPY and n = 9 VIP).

**Table 2.** The latency and jitter of first elicited oEPSPs and oEPSCs were low in both NYP and VIP neurons. Values show mean ± standard error of the mean. NPY: n = 24, N = 15, VIP: n = 26, N = 17.

|  | I Clamp<br>NPY EPSP | I Clamp<br>VIP EPSP | V Clamp<br>NPY EPSC | V Clamp<br>VIP EPSC |
| --- | --- | --- | --- | --- |
| **Latency (ms)** | 2.05 ± 0.2 | 2.42 ± 0.1 | 1.57 ± 0.1 | 1.98 ± 0.17 |
| **Jitter (ms)** | 0.35 ± 0.1 | 0.73 ± 0.1 | 0.26 ± 0.04 | 0.28 ± 0.04 |

Interestingly, we observed several instances in which optically stimulating T-stellate axons elicited a short-latency excitatory response followed by one or more longer-latency inhibitory (**Figure 5B**) or excitatory responses (**Figure 5C**). Because the first event in these complex responses had a short latency and low jitter, it likely represented monosynaptic input from T-stellate terminals (inhibition: NPY, latency = 1.92 ± 0.2 ms, jitter = 0.17 ± 0.06 ms (n = 6, N = 4); VIP, latency = 1.77 ± 0.08 ms, jitter = 0.08 ± 0.04 ms (n = 6, N = 5); excitation: NPY, latency = 1.58 ± 0.10 ms, jitter = 0.27 ± 0.04 ms (n = 19, N = 11); VIP, latency = 2.05 ± 0.30 ms, jitter = 0.41 ± 0.07 ms (n = 9, N = 9); compare to **Table 2**). The subsequent responses presumably resulted from polysynaptic pathways in which optogenetically activated T-stellate synapses elicited firing in excitatory or inhibitory neurons that in turn provided feedforward synaptic input to the recorded neuron. Since these experiments were performed in IC brain slices, the feedforward neurons were almost certainly other IC neurons. Together, these data show that T-stellate neurons provide direct input to many NPY and VIP neurons in the ICc, and that T-stellate projections can recruit inhibitory and excitatory feedforward circuits in the IC.

### T-stellate input to ICc neurons undergoes short-term synaptic depression

To determine if T-stellate synapses onto NPY and VIP neurons exhibit short-term synaptic plasticity, we delivered trains of 10 optogenetic stimuli at 20, 50, and 70 Hz to repetitively activate Chronos-EGFP-expressing T-stellate projections while making voltage-clamp recordings from NPY and VIP neurons in the ICc. Light pulse intensity was set using a minimum stimulation paradigm in which most, but not all, light presentations elicited an oEPSC. All metrics reported include only successful oEPSCs and not failures. First, we compared the amplitude of the first oEPSCs elicited between NPY and VIP neurons. For this analysis, we also included data from earlier experiments where we were optimizing different train parameters, as varying train parameters did not affect the first oEPSCs elicited. We found that, similar to our current clamp experiments, the first oEPSCs elicited in NPY neurons were significantly larger than those in VIP neurons (NPY = −94.8 ± 10.4 pA (n = 24, N = 15), VIP = −47.9 ± 5.14 pA (n = 26, N = 17), t = −4.1, df = 543.2, *p* < 0.001, Welch’s t-test). Next, we found that 20 Hz trains of 10 light pulses elicited oEPSCs that significantly decreased in amplitude as a function of light flash number for both NPY and VIP neurons (NPY: LMM, main effect of light pulse number, *F*_9,63_ = 15.221, *p* < 0.001 (n = 8, N = 4); VIP: LMM, main effect of light pulse number, *F*_9,72_ = 15.247, *p* < 0.001, (n = 9, N = 5); statistics for pairwise comparisons of oEPSCs_2-10_ to oEPSC_1_ are shown in **Supplemental Table 1**), indicating that these synapses undergo short-term depression (STD) (**Figure 6A-B**). However, the magnitude of STD elicited by 20 Hz stimulation did not significantly differ between NPY and VIP neurons (LMM, main effect of neuron type, *F*_1,15_ = 0.072, *p* = 0.792) (**Figure 6C**). We observed similar trends with 50 Hz and 70 Hz stimulation: the magnitude of oEPSCs significantly decreased with increasing light flash number (50 Hz NPY: LMM, main effect of light pulse number, *F*_9,63_ = 24.089, *p* < 0.001 (n = 8, N = 4); 50 Hz VIP: LMM, main effect of light pulse number, *F*_9,72_ = 21.932, *p* < 0.001 (n = 9, N = 5); 70 Hz NPY: LMM, *F*_9,36_ = 15.868, *p* < 0.001, (n = 5, N = 4); 70 Hz VIP: LMM, *F*_9,54_ = 23.534, p < 0.001, (n = 7, N = 5); statistics for pairwise comparisons of oEPSCs_2-10_ to oEPSC_1_ are shown in **Supplemental Table 1**), but there was no significant difference in STD between NPY and VIP neurons (50 Hz: LMM, main effect of neuron type, *F*_1,15_ = 0.194, *p* = 0.666; 70Hz: LMM, main effect of neuron type, *F*_1,10_ = 0.001, *p* = 0.917) (**Figure 6D-I**). We also found that the STD at both NPY and VIP neurons significantly changed with frequency of activation (NPY: LMM, main effect of frequency, *F*_2,192.01_ = 39.878, *p* < 0.001; VIP: LMM, main effect of frequency, *F*_2,230.41_ = 24.648, *p* < 0.001; statistics for pairwise comparisons across frequencies are shown in **Supplemental Table 6**) (**Figure 6J**). Together, these data show that T-stellate synapses on NPY and VIP neurons exhibit STD that increases at higher stimulation frequencies and that the extent of STD was similar at synapses onto both neuron types.

**Figure 6.**
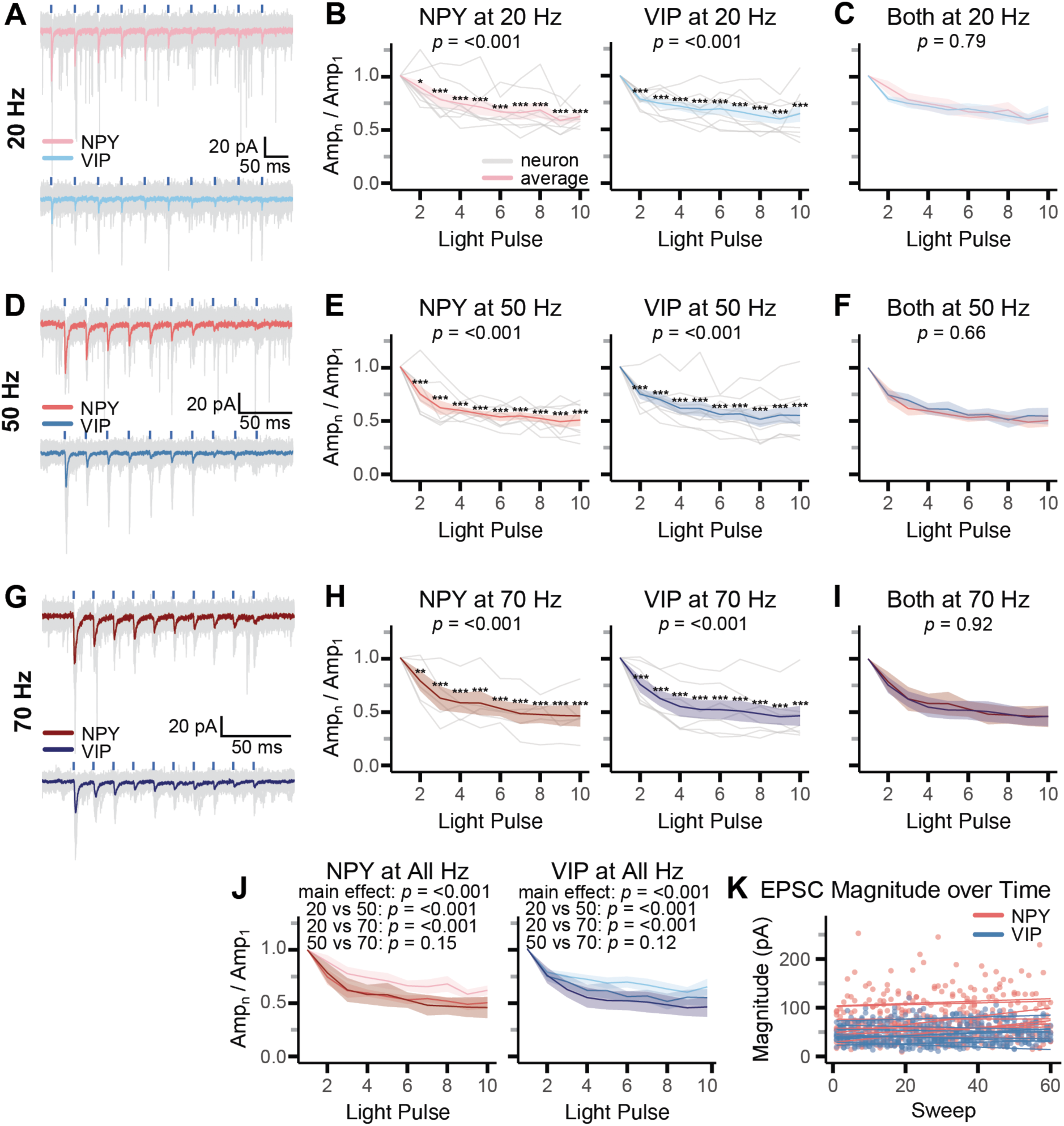
T-stellate input to NPY and VIP neurons undergoes short-term synaptic depression (STD). **A.** 20 Hz trains of light pulses elicited trains of oEPSCs in a representative NPY neuron (top) and a representative VIP neuron (bottom). Traces from individual trials are shown in gray, and average responses are overlaid in color. Light pulses are indicated by blue bars. **B.** T-stellate input to both NPY and VIP neurons exhibited STD with 20 Hz stimulation. **C**. 20 Hz stimulation elicited similar amounts of STD at T-stellate synapses onto NPY neurons and VIP neurons. Average responses are shown. **D-I.** 50 Hz stimulus trains (**D-F**) and 70 Hz stimulus trains (**G-I**) also elicited STD at T-stellate synapses onto NPY and VIP neurons. Data are presented as in A-C. **J.** In both NPY and VIP neurons, the amount of STD elicited by 50 Hz and 70 Hz stimulus trains was greater than that elicited by 20 Hz trains. **K.** The amplitude of the first oEPSCs evoked by stimulus trains did not change significantly across trials suggesting that the oEPSC amplitude changes during trials were due to STD and not recording rundown. Lines represent linear fits to first oEPSC amplitudes from each train for an individual cell. In **B,C,E,F,H-J**, data from individual cells are shown in gray, and averages across cells are overlaid in color. Shaded regions around average lines indicate SEM. *p*-values for main effects, as determined by LMM tests, are indicated at the top of plots. Asterisks indicate significant differences determined by post hoc tests comparing the response at specific light pulse numbers to the response elicited by the first light pulse in the train. * = *p* < 0.05, ** = *p* < 0.01, *** = *p* < 0.001. In **J,** additional *p*-values indicate post hoc pairwise comparisons of group data from different train frequencies.

To check that we were allowing sufficient time for recovery from STD between delivering trains of stimuli and that repeated optogenetic stimulation did not lead to run down in the strength of T-stellate input over time, we assessed the amplitude of the first oEPSC in each train as a function of train presentation number. We then fit a line to those first oEPSC amplitudes and found that, for all cells, the slope of the line was nearly zero, and the slopes did not significantly differ between NPY and VIP neurons (NPY mean slope: 0.29 ± 0.34 pA/train presentation; VIP mean slope: 0.01 ± 0.23 pA/train presentation; t = 1.97, df = 11.84, *p* = 0.07, Welch’s t-test) (**Figure 6K**).

### Changing AAV serotype and optical stimulus location did not alter STD at T-stellate synapses

An important consideration when using optogenetics to investigate short-term synaptic plasticity is that the AAV serotype and the location of light activation can result in STD at synapses between some, but not all, neuron types that are otherwise known to exhibit short-term potentiation when electrically stimulated (Zhang & Oertner, 2007; Klapoetke *et al*., 2014; Jackman *et al*., 2014). Throughout the present study, we used AAV1 vectors to drive Chronos-GFP expression in T-stellate neurons because this serotype yielded high expression of Chronos-GFP in T-stellate projections, which enhanced the chances of finding NPY or VIP neurons in the IC that received T-stellate input. Similarly, we delivered light pulses through the microscope objective with the recorded cell centered in the field of view to enhance the probability of activating T-stellate synapses onto NPY or VIP neurons. This mode of optical stimulation has been dubbed “on-bouton” stimulation because it can elicit vesicular release in two ways: 1) by activating channelrhodopsins expressed in synaptic terminals, driving depolarization and vesicular release in an action-potential independent way, and 2) by activating channelrhodopsins in axons, leading to action potentials that in turn invade synaptic terminals to elicit vesicular release. It was therefore important to test whether the STD we observed was an artifact of the AAV1 serotype or the illumination pattern used.

Since Jackman and colleagues showed that transduction with AAV9 vectors did not lead to spurious STD, we injected AAV9-hSyn-Chronos-EGFP into the VCN of NPY mice. After allowing several weeks for Chronos-EGFP expression, we prepared IC slices and measured short-term plasticity using on-bouton activation, where the light pulses were delivered over the recorded neuron, and off-bouton activation, where the light pulses were delivered ∼430 µm away from the recorded neuron. We found that the AAV9 serotype had a lower transduction efficiency in the VCN, which made it more difficult to find NPY neurons that received sufficiently large T-stellate input to examine short-term plasticity. Because our earlier experiments showed that T-stellate input elicits smaller oEPSCs in VIP neurons than NPY neurons and the effects of AAV transduction are presynaptic (Jackman *et al*., 2014), we focused on NPY neurons for this experiment.

In recordings from NPY neurons, T-stellate synapses continued to exhibit significant STD when the AAV9 vector was used, whether activated with on-bouton or off-bouton light pulses (on-bouton: LMM, main effect of light pulse number, *F*_9,63_ = 4.96, *p* < 0.001; off-bouton: LMM, main effect of light pulse number, *F*_9,63_ = 4.13, *p* < 0.001, (n = 8, N = 6); statistics for pairwise comparisons of oEPSCs_2-10_ to oEPSC_1_ are shown in **Supplemental Table 2**) (**Figure 7A**). The magnitude of STD also did not differ between on-bouton and off-bouton stimulation (LMM, main effect of activation location, *F*_1,14_ = 0.07, *p* = 0.789) (**Figure 7B**). We next compared the magnitude of STD between AAV1 on-bouton and AAV9 on-bouton activation and found no significant difference between groups (LMM, main effect of AAV serotype, *F*_1,142_ = 1.95, *p* = 0.164) (**Figure 7C**). Furthermore, there was no difference between AAV1 on-bouton and AAV9 off-bouton activation (LMM, main effect, *F*_1,14_ = 0.790, *p* = 0.389) (**Figure 7D**). Together, these data suggest that T-stellate projections to the IC are not susceptible to the spurious STD that AAV1 transduction and on-bouton stimulation can cause at some synapses.

**Figure 7.**
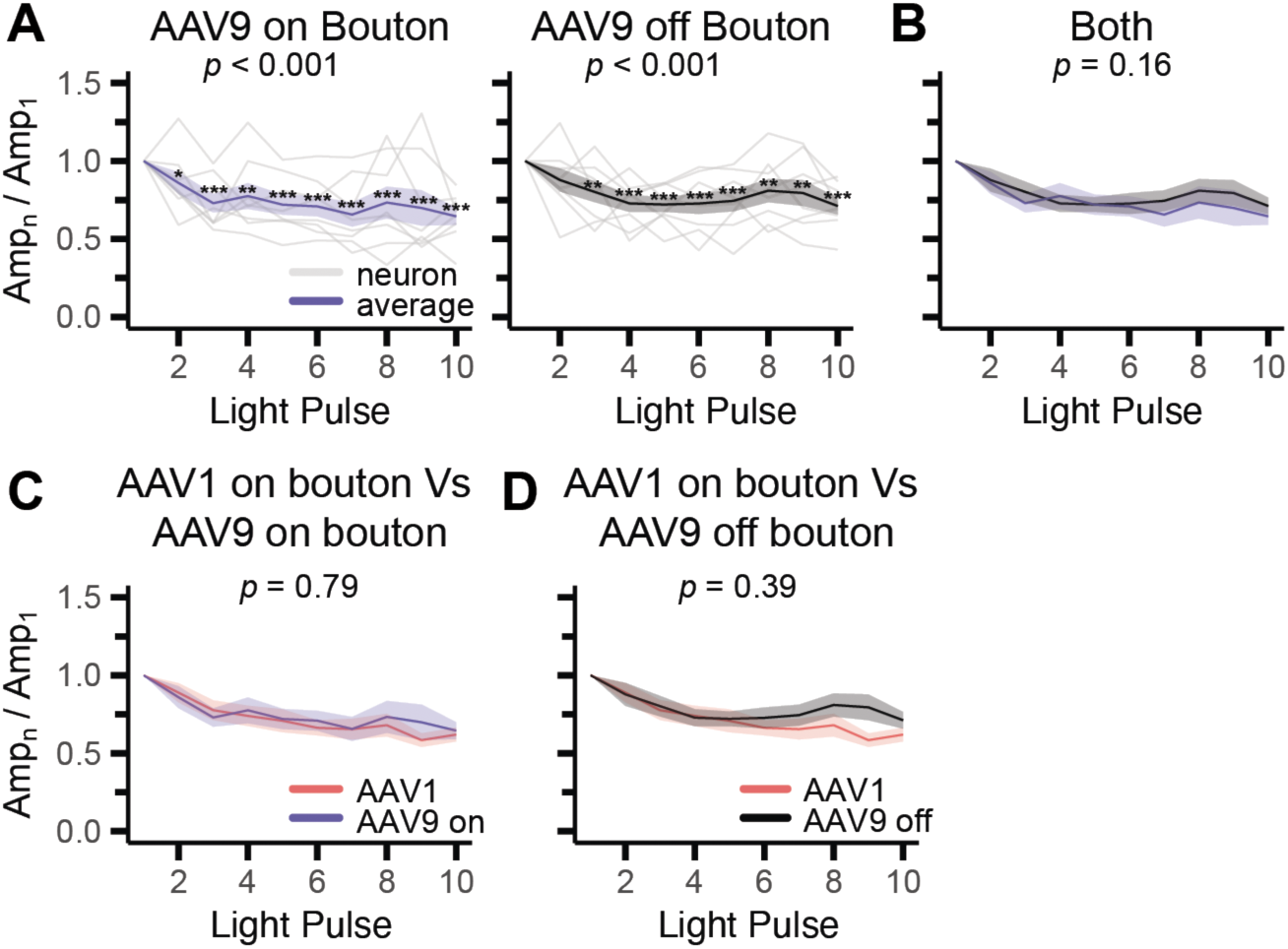
T-stellate synapses exhibited similar patterns of short-term plasticity when transduced with AAV9 and activated with off-bouton stimulation. **A.** In recordings from NPY neurons, T-stellate projections transduced with AAV9-hSyn-Chronos-EGFP exhibit STD in response to 20 Hz activation both with on-bouton and off-bouton activation. **B.** There was no significant difference between the STD elicited with on-bouton and off-bouton optical stimulation. **C.** AAV serotype did not alter the STD exhibited at T-stellate synapses onto NPY neurons when on-bouton stimulation was used. **D.** Similarly, there was no significant difference in the STD exhibited by AAV1-transduced T-stellate projections activated with on-bouton stimulation and that exhibited by AAV9-transduced T-stellate projections activated with off-bouton stimulation. In **A-D,** data from individual cells are shown in gray and averages across cells are overlaid in color. Shaded regions around average lines indicate SEM. The *p*-values for main effects, as determined by LMM tests, are indicated at the top of plots. Asterisks indicate significant differences determined by post hoc tests comparing the response at specific light pulse numbers to the response elicited by the first light pulse in the train. * = *p* < 0.05, ** = *p* < 0.01, *** = *p* < 0.001.

### oEPSC failure rate and latency increased during trains

We next examined the reliability and temporal precision of oEPSCs evoked with train stimuli. Due to the minimal stimulation paradigm we used, not all light pulses elicited a postsynaptic response. With 20 Hz activation, oEPSC failure rate significantly increased with light pulse number at both NPY and VIP neurons (NPY: LMM, main effect of light pulse number, *F*_9,63_ = 4.44, *p* < 0.001; VIP: LMM, main effect of light pulse number, *F*_9,72_ = 7.64, *p* < 0.001; statistics for pairwise comparisons of oEPSCs_2-10_ to oEPSC_1_ are shown in **Supplemental Table 3**), but there was no significant difference in failure rate between the two neuron types (LMM, main effect of neuron type, *F*_1,15_ = 0.60, *p* = 0.449) (**Figure 8A-B**). We also observed similar trends with 50 and 70 Hz trains; the failure rate significantly increased with light pulse number (50 Hz NPY: LMM, main effect of light pulse number, *F*_9,63_ = 3.37, *p* < 0.001; 50 Hz VIP: LMM, main effect of light pulse number, *F*_9,72_ = 14.55, *p* < 0.001; 70 Hz NPY: LMM, *F*_9,36_ = 2.97, *p* < 0.001; 70 Hz VIP: LMM, *F*_9,54_ = 5.53, p < 0.001; statistics for pairwise comparisons of oEPSCs_2-10_ to oEPSC_1_ are shown in **Supplemental Table 3**), but there was no difference in failure rate between NPY and VIP neurons (50 Hz: LMM, main effect of neuron type, *F*_1,15_ = 0.650, *p* = 0.433; 70Hz: LMM, *F*_1,10_ = 0.044, *p* = 0.838) (**Figure 8C-F**). For both NPY and VIP neurons, the failure rate was not significantly different across stimulation frequencies (NPY: LMM, main effect of frequency, *F*_2,192.01_ = 1.022, *p* = 0.362; VIP: LMM, main effect of frequency, *F*_2,230.29_ = 1.49, *p* = 0.227; statistics for pairwise comparisons across frequencies are shown in **Supplemental Table 6**) (**Figure 8G**).

**Figure 8.**
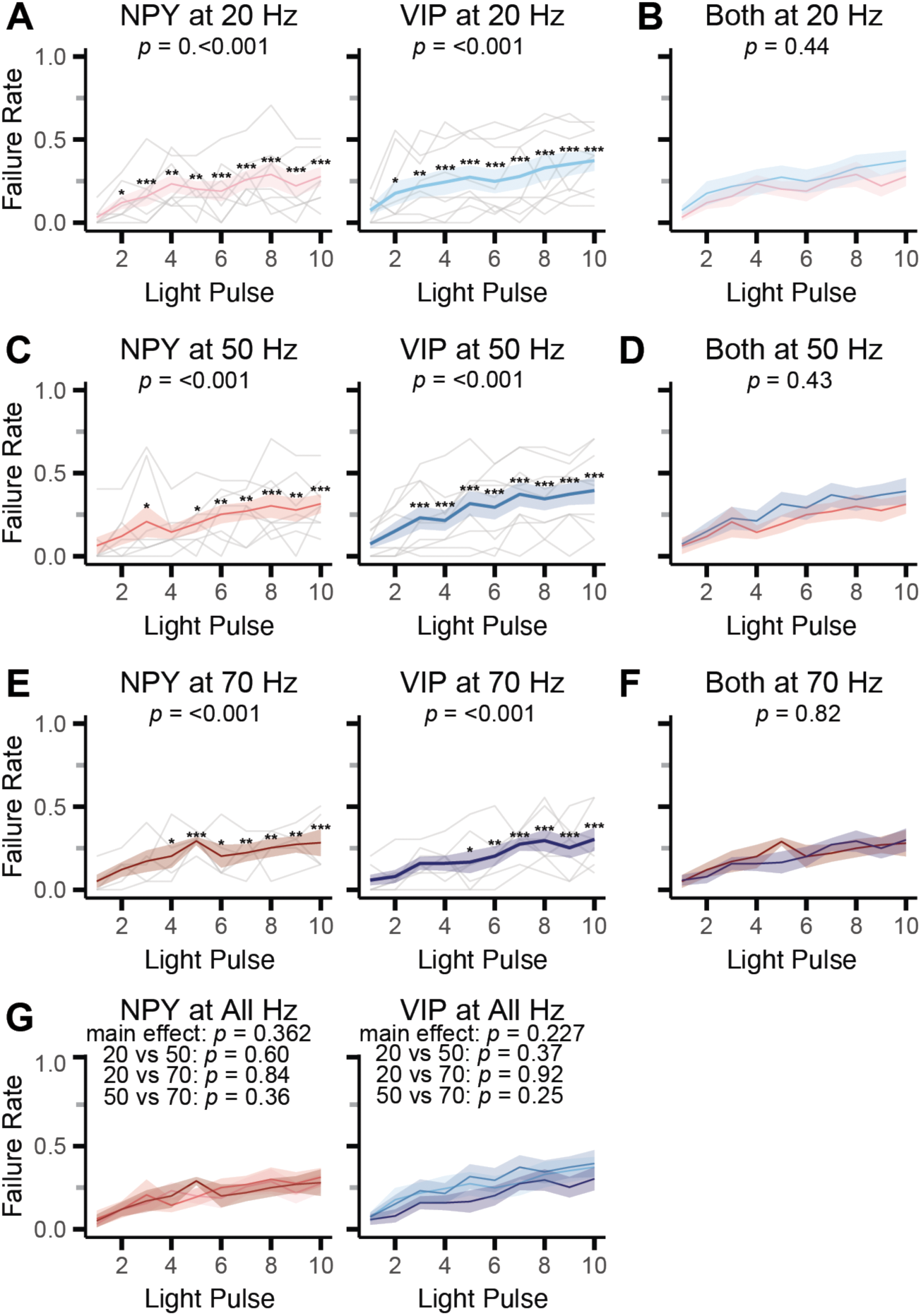
oEPSC failure rate increased during train stimulation. **A.** oEPSC failure rate increased during trains in both NPY neurons (left) and VIP neurons (right). **B.** There was no significant difference in oEPSC failure rates between NPY and VIP neurons. **C-F.** oEPSC failure rates also increased during 50 Hz trains (**C**) and 70 Hz trains (**E**), but the failure rates did not significantly differ between NPY and VIP neurons (**D,F**). Data presented as in **A,B**. **G.** oEPSC failure rates were not significantly different across stimulation frequencies. In **A-G**, data from individual cells are shown in gray, and averages across cells are overlaid in color. Shaded regions around average lines indicate SEM. The *p*-values for main effects, as determined by LMM tests, are indicated at the top of plots. Asterisks indicate significant differences determined by post hoc tests comparing responses at specific light pulse numbers to the response elicited by the first light pulse in the train. * = *p* < 0.05, ** = *p* < 0.01, *** = *p* < 0.001. In **G,** additional *p*-values indicate post hoc pairwise comparisons of group data from different train frequencies.

We quantified the latency of each oEPSC as a function of light pulse number for all train frequencies by measuring the time from light pulse onset to the 10% rise of the oEPSC. We found that with 20 Hz trains, oEPSC latency significantly increased with light pulse number in both NPY and VIP neurons (NPY: LMM, main effect of light pulse number, *F*_9,63_ = 6.490, *p* < 0.001; VIP: LMM, main effect of light pulse number, *F*_9,72_ = 8.337, *p* < 0.001; statistics for pairwise comparisons of oEPSCs_2-10_ to oEPSC_1_ are shown in **Supplemental Table 4**), but the latency was not significantly different between NPY and VIP neurons (LMM, main effect of neuron type, *F*_1,15_ = 0.464, *p* = 0.506) (**Figure 9A-B**). With 50 Hz trains, oEPSC latency also increased with light pulse number (NPY: LMM, main effect of light pulse number, *F*_9,63_ = 5.298, *p* < 0.001; VIP: LMM, main effect of light pulse number, *F*_9,72_ = 12.14, *p* < 0.001; statistics for pairwise comparisons of oEPSCs_2-10_ to oEPSC_1_ are shown in **Supplemental Table 4**), but did not differ between neuron types (LMM, main effect of neuron type, *F*_1,15_ = 0.349, *p* = 0.564) (**Figure 9C**). With 70 Hz trains, oEPSC latency significantly increased with light pulse number only in VIP neurons (NPY: LMM, main effect of light pulse number, *F*_9,36_ = 1.999, *p* = 0.068; VIP: LMM, main effect of light pulse number, *F*_9,54_ = 5.207, *p* < 0.001; statistics for pairwise comparisons of oEPSCs_2-10_ to oEPSC_1_ are shown in **Supplemental Table 4**) (**Figure 9E**), but there was no significant difference in the latency between cell types (LMM, main effect of neuron type, *F*_1,10_ = 1.250, *p* = 0.290). oEPSC latency during train stimuli did not significantly differ as a function of stimulation train frequency in NPY neurons but did differ in VIP neurons (NPY: LMM, main effect of frequency, *F*_2,192.01_ = 1.022, *p* = 0.082; VIP: LMM, main effect of frequency, *F*_2,230.29_ = 19.115, *p* < 0.001; statistics for pairwise comparisons across frequencies are shown in **Supplemental Table 6) (Figure 9G).**

**Figure 9.**
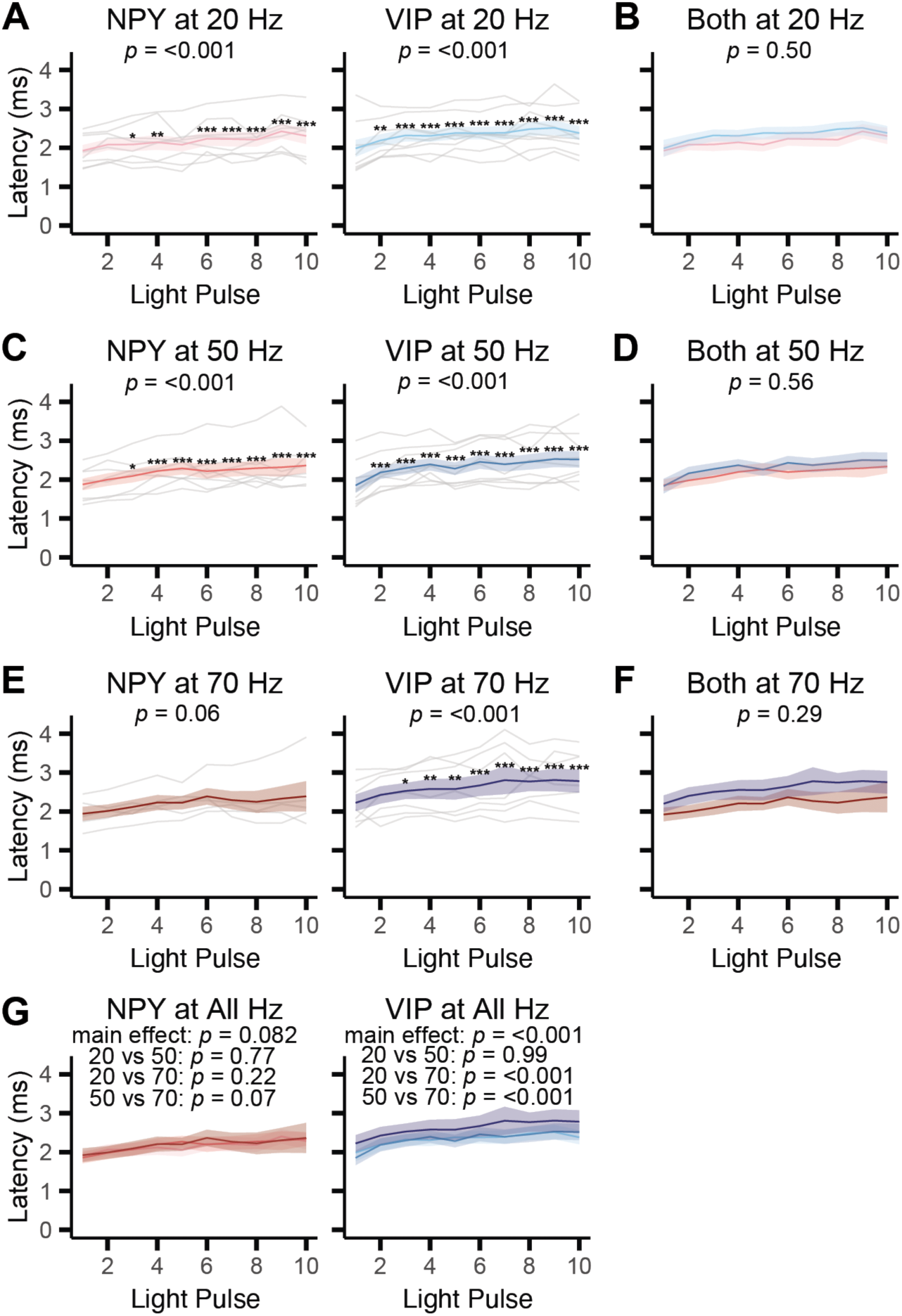
oEPSC latency increased during train stimulation. **A.** The latency from light pulse onset to the 10% rise of oEPSCs increased during trains in both NPY neurons (left) and VIP neurons (right). **B.** There was no significant difference in oEPSC latencies between NPY and VIP neurons. **C-F.** oEPSC latencies also increased during 50 Hz trains (**C**) and 70 Hz trains (**E**), but latencies did not significantly differ between NPY and VIP neurons (**D,F**). Data presented as in **A,B**. **G.** In NPY neurons, oEPSC latencies did not significantly differ across stimulation frequencies, but in VIP neurons oEPSC latency was higher in response to 70 Hz trains compared to 20 Hz and 50 Hz trains. In **A-G**, data from individual cells are shown in gray, and averages across cells are overlaid in color. Shaded regions around average lines indicate SEM. The *p*-values for main effects, as determined by LMM tests, are indicated at the top of plots. Asterisks indicate significant differences determined by post hoc tests comparing the response at specific light pulse numbers to the response elicited by the first light pulse in the train. * = *p* < 0.05, ** = *p* < 0.01, *** = *p* < 0.001. In **G,** additional *p*-values indicate post hoc pairwise comparisons of group data from different train frequencies.

Finally, we measured the jitter in oEPSC latencies by calculating the standard deviation of latencies across trials. We found that jitter significantly increased across light pulse trains of all frequencies in VIP neurons, but not NPY neurons (20 Hz NPY: LMM, main effect of light pulse number, *F*_9,63_ = 1.697, *p* = 0.108; 20 Hz VIP: LMM, main effect of light pulse number, *F*_9,72_ = 2.077, *p* = 0.043; 50 Hz NPY: LMM, *F*_9,63_ = 1.321, *p* = 0.244; 50 Hz VIP: LMM, *F*_9,72_ = 2.246, *p* = 0.028; 70 Hz NPY: LMM, *F*_9,36_ = 0.958, *p* = 0.490; 70 Hz VIP: LMM, *F*_9,54_ = 2.761, p = 0.010; statistics for pairwise comparisons of oEPSCs_2-10_ to oEPSC_1_ are shown in **Supplemental Table 5**) (**Figure 10A,C, E**). However, oEPSC jitter did not significantly differ between NPY and VIP neurons across all activation frequencies (20 Hz: LMM, main effect of neuron type, *F*_1,15_ = 0.181, *p* = 0.676; 50 Hz: LMM, *F*_1,15_ = 1.853, *p* = 0.194; 70Hz: LMM, *F*_1,10_ = 2.308, *p* = 0.160) (**Figure 10B,D, F**). We also found that the jitter at both NPY and VIP neurons significantly changed with frequency of activation (NPY: LMM, main effect of frequency, *F*_2,192.01_ = 8.995, *p* < 0.001; VIP: LMM, main effect of frequency, *F*_2,233.59_ = 3.343, *p* = 0.014; statistics for pairwise comparisons across frequencies are shown in **Supplemental Table 6**) (**Figure 10G**). Together, these results indicate that the reliability and temporal precision of optically evoked T-stellate input to NPY and VIP neurons decreased during stimulus trains, albeit with modest effect sizes. Since we also found that T-stellate synapses exhibit STD, our data suggest that synaptic input from T-stellate neurons to NPY and VIP neurons decreases in strength, reliability, and temporal precision during periods of sustained T-stellate neuron activity.

**Figure 10.**
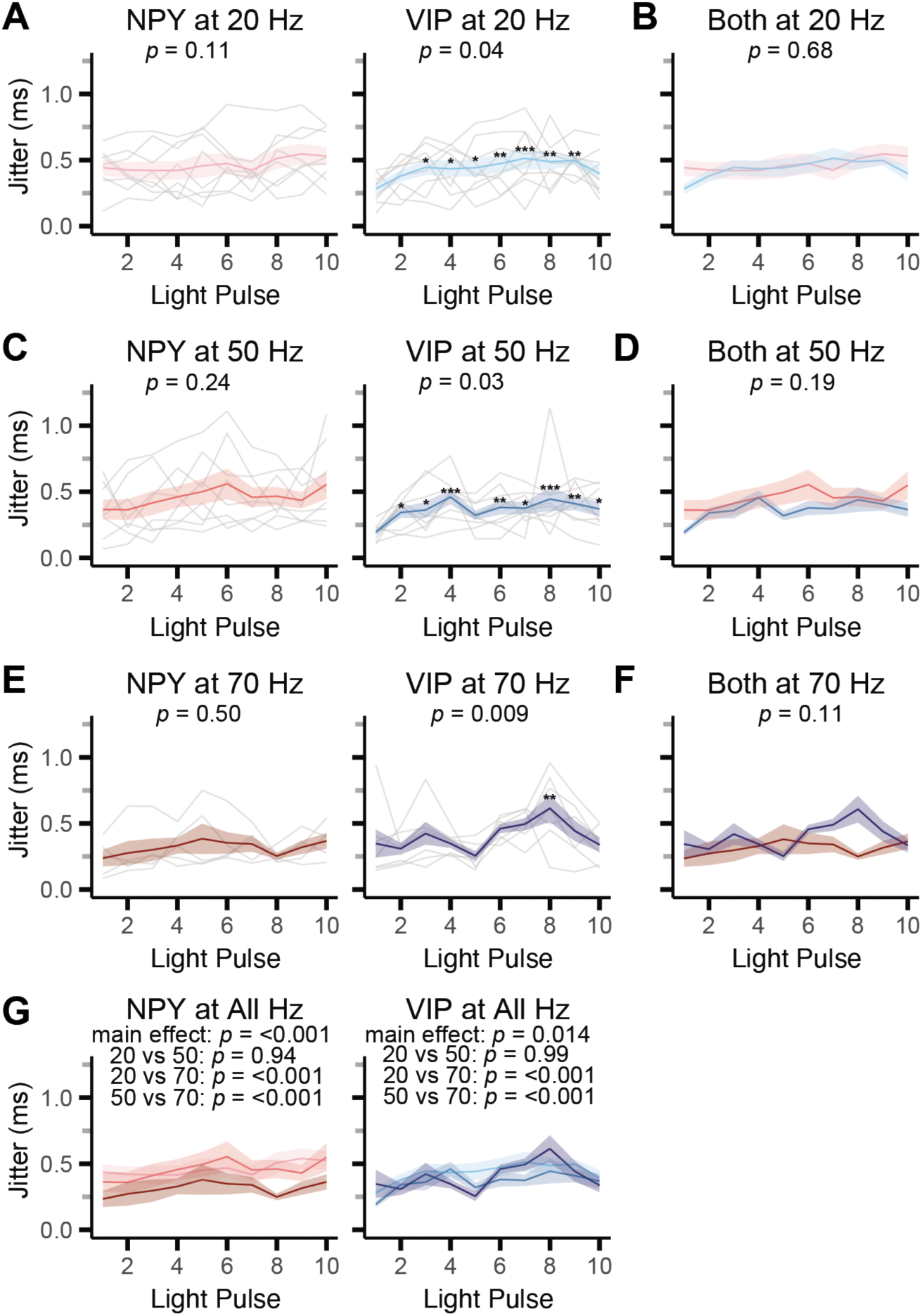
Jitter in the latency of oEPSCs did not change during stimulus trains for NPY neurons but increased slightly for VIP neurons. **A.** oEPSC jitter was measured as the standard deviation of oEPSC latency. Jitter did not change significantly during 20 Hz stimulus trains in NPY neurons (left) but slightly increased in VIP neurons (right). **B.** oEPSC jitter did not significantly differ between NPY and VIP neurons. **C-F.** Similarly, oEPSC jitter did not significantly change during 50 Hz (**C**) and 70 Hz (**E**) stimulus trains in NPY neurons but slightly increased in VIP neurons. However, there was no significant difference in oEPSC jitter between NPY and VIP neurons at either stimulus frequency (**D,F**). **G.** In both NPY and VIP neurons, oEPSC jitter was significantly different in response to 70 Hz stimulus trains compared to 20 Hz and 50 Hz trains. In **A-G**, data from individual cells are shown in gray, and averages across cells are overlaid in color. Shaded regions around average lines indicate SEM. The *p*-values for group effects, as determined by LMM tests, are indicated at the top of plots. Asterisks indicate significant differences determined by post hoc tests comparing responses at specific light pulse numbers to the response elicited by the first light pulse in the train. * = *p* < 0.05, ** = *p* < 0.01, *** = *p* < 0.001. In **G,** additional *p*-values indicate post hoc pairwise comparisons of group data from different train frequencies.

## Discussion

We used channelrhodopsin assisted circuit mapping to assess the prevalence, amplitude, kinetics, and short-term synaptic plasticity of T-stellate input from the VCN to two molecularly identified neuron types in the IC. We found that both inhibitory NPY neurons and excitatory VIP neurons receive excitatory synaptic input from T-stellate neurons. In addition, activation of T-stellate projections drove local sources of feedforward and/or feedback input to NPY and VIP neurons. These results demonstrate that a single ascending source of auditory input can engage both inhibitory and excitatory IC neurons and elicit complex local circuit operations in the IC. Our recordings also showed that T-stellate input was more prevalent and evoked larger and faster postsynaptic responses in NPY neurons than VIP neurons, but T-stellate synapses onto both NPY and VIP neurons exhibited similar patterns of STD in response to stimulus trains. The extent of STD at T-stellate synapses increased with stimulation frequency, suggesting that T-stellate input to the IC adapts in a frequency-dependent way during periods of ongoing activity. Taken together, our results provide mechanistic insights into how ascending inputs interface with IC circuits to shape auditory computations.

### Implications of connections between T-stellate projections and IC circuits

The IC is a major integrative hub for ascending and descending pathways in the central auditory system, but it is not known whether individual pathways preferentially contact specific IC neuron types. Previous anatomical studies showed that auditory brain regions projecting to the IC can contact both excitatory and inhibitory IC neurons (Chen *et al*., 2018). Similarly, cholinergic projections from the pedunculopontine tegmental nucleus contact both glutamatergic and GABAergic IC neurons (Noftz *et al*., 2020). These studies suggest that inputs to the IC are divergent, contacting at least two postsynaptic populations of neurons. Our results provide a new level of insight by providing functional and anatomical evidence that T-stellate projections contact at least two molecularly defined neuron types in the IC: inhibitory NPY neurons and excitatory VIP neurons. Given the size of the T-stellate projection to the IC (Ryugo & Milinkeviciute, 2023) and the fact that NPY and VIP neurons account for only a portion of IC neurons (Goyer *et al*., 2019; Silveira *et al*., 2020; Beebe *et al*., 2022; Silveira *et al*., 2024), it is likely that T-stellate projections also contact other neuron types in the IC. As molecular markers for additional IC neuron types are discovered (Drotos & Roberts, 2024), it will become possible to determine the full complement of IC neuron types that T-stellate projections contact, a key step to understanding how the acoustic information carried by T-stellate neurons is processed in the IC.

The finding that T-stellate neurons synapse onto both NPY and VIP neurons suggests that T-stellate input can recruit complex local circuit operations in the IC. For example, since both NPY and VIP neurons have local axon collaterals (Goyer *et al*., 2019; Beebe *et al*., 2022; Silveira *et al*., 2024), T-stellate input may activate feedforward inhibitory and feedback excitatory circuits in the IC. Indeed, our recordings showed that activating T-stellate projections routinely elicited feedforward (polysynaptic) IPSPs and EPSCs (**Figure 5B,C**). Among its many roles, feedforward inhibition can limit the time window for synaptic integration (Pouille & Scanziani, 2001) and regulate the dynamic range of postsynaptic neurons (Pouille *et al*., 2009). Feedback excitation gives rise to recurrent excitation, a prominent feature of IC local circuits (Oberle *et al*., 2023; Silveira *et al*., 2023). In cerebral cortex, recurrent excitation can refine sensory receptive fields by amplifying, suppressing, and/or filtering specific stimulus features (Lien & Scanziani, 2013; Li *et al*., 2013a, 2013b; Niell & Scanziani, 2021; Oldenburg *et al*., 2024; Deveau *et al*., 2026). An important goal for future studies will be to determine whether T-stellate projections recruit similar local circuit operations in the IC.

### Technical limitations

Optogenetics has revolutionized the study of synaptic connections by enabling experimenters to selectively manipulate anatomically and/or molecularly targeted populations of neurons (Boyden *et al*., 2005; Petreanu *et al*., 2007, 2009), but some experimental limitations must be considered. In particular, Jackman and colleagues found that using AAV1 serotype viruses to express channelrhodopsin-2 (ChR2) led to artificial STD at synapses between some neuron types but not others (Jackman *et al*., 2014). The reasons for this remain unclear, but artificial STD did not occur when AAV9 serotype viruses were used (Jackman *et al*., 2014). In the present study, we primarily relied on AAV1 viruses because AAV1 drove higher Chronos expression than other AAV serotypes, increasing the probability that we would find IC neurons receiving optically evoked T-stellate input and reducing the animal numbers needed. We therefore conducted control experiments using AAV9 to check for artificial STD, but we found no difference between AAV1 and AAV9 in the amount of STD observed at T-stellate synapses (**Figure 7C**).

The site of optical stimulation can also affect short-term plasticity (Jackman *et al*., 2014), but we saw no difference in the amount of STD elicited by on-bouton stimulation of T-stellate projections transduced with AAV1 and off-bouton stimulation of T-stellate projections transduced with AAV9 (**Figure 7D**). Thus, our data suggest that the STD we observed at T-stellate synapses in the IC is a genuine synaptic feature and not attributable to artificial STD. This is similar to the situation for Purkinje cell synapses onto deep cerebellar nuclei neurons, which Jackman and colleagues found were also not sensitive to AAV serotype or the location of optical stimulation (Jackman *et al*., 2014).

A final consideration is that T-stellate neurons can fire at frequencies up to several hundred Hz (Rhode & Smith, 1986; Young *et al*., 1988; Blackburn & Sachs, 1989), but it is not possible stimulate at such high rates with currently available optogenetic tools. We used Chronos here because its kinetics are among the fastest of the currently available channelrhodopsins (Klapoetke *et al*., 2014; Keppeler *et al*., 2018), but in our hands, T-stellate projections could not be reliably driven by Chronos at frequencies higher than 70 Hz. It therefore remains unclear whether STD at T-stellate synapses onto NPY and VIP neurons continues to increase at higher stimulation frequencies or if the similar amount of STD elicited by 50 Hz and 70 Hz stimulation trains (**Figure 6J**) indicates that STD at T-stellate synapses reaches a stable level above 50 Hz. Electrical stimulation is an appealing alternative, as it would allow for higher stimulation rates and is not subject to artificial STD. However, electrical stimulation is not a viable option for selectively stimulating T-stellate projections in IC brain slices because the axons of T-stellate neurons ascend through the lemniscal tract, where they mix with numerous other sources of ascending input to the IC (Adams, 1979; Oliver, 1987; Budinger *et al*., 2000; Malmierca *et al*., 2005). Thus, with appropriate controls, optogenetic stimulation is a valuable approach for investigating the synaptic physiology of defined sources of input to the IC.

### Implications of STD at T-stellate synapses for auditory processing

Previous studies highlight a variety of roles for short-term plasticity in sensory adaptation (Chung *et al*., 2002; Brecht & Sakmann, 2002; Katz *et al*., 2006; Heiss *et al*., 2008; Holla *et al*., 2026), implementing temporal filters (Buonomano & Merzenich, 1995; Fortune & Rose, 2000; Rosenbaum *et al*., 2012; Mondal *et al*., 2022; Barri *et al*., 2022), and gain control (Abbott *et al*., 1997; Rothman *et al*., 2009). Here, we focus on two mechanisms that seem particularly relevant to the IC. First, STD of afferent synapses is a compelling and simple mechanism for sensory adaptation. For example, retinogeniculate synapses exhibit pronounced STD in vitro (Turner & Salt, 1998), and repeated whisker stimulation leads to STD of thalamocortical synapses in rodent somatosensory cortex (Chung *et al*., 2002; Brecht & Sakmann, 2002; Katz *et al*., 2006). Auditory nerve fibers also exhibit STD at their synapses in the VCN, but interestingly, they exhibit less STD onto T-stellate neurons than other VCN neuron types (Wu & Oertel, 1987; Chanda & Xu-Friedman, 2010; Cao & Oertel, 2010). Our results suggest that this may be compensated for by STD at T-stellate synapses in the IC. Straightforward interpretations like this, however, are confounded by studies showing that, under awake in vivo conditions, afferent synapses spend much of the time in an already depressed state, likely due to higher ongoing activity levels in the awake brain (Boudreau & Ferster, 2005; Reig *et al*., 2006; Hermann *et al*., 2007; Weyand, 2007; Lorteije *et al*., 2009).

Second, several studies have shown that short-term plasticity can interact with intrinsic neurophysiology and circuit architecture in complex ways that give rise to more nuanced computations (Abbott *et al*., 1997; Heiss *et al*., 2008; Rothman *et al*., 2009). In a foundational study, Fortune and Rose examined temporal filtering in midbrain electrosensory neurons in the torus semicircularis of weakly electric fish (Fortune & Rose, 2000). They found that STD at ascending synapses implemented a low-pass filter that reduced the influence of repeated sensory signals, such as those representing interactions with the electric organ discharges of nearby weakly electric fish. Intriguingly, short bursts of higher-frequency input, representing the appearance of salient objects in the neighboring environment, were enhanced due to short-term facilitation. Since the torus semicircularis and the IC are homologous structures (Foster & Hall, 1978), short-term plasticity may similarly enhance the capacity of IC circuits to preferentially detect rapid changes in sensory input. A recent study from Holla and colleagues showed that cerebellar circuits implement a conceptually similar computation. Using in vivo recordings and stimulation of the whisker pads of mice, the authors showed that short-term plasticity interacts with cerebellar circuits to enhance the detection of changes in sensory input while dampening responses to repeated sensory stimuli and sensory-evoked whisker movement (Holla *et al*., 2026).

By analogy to these studies, it is tempting to speculate that STD at T-stellate synapses in the IC enhances the ability of IC circuits to detect rapid changes in acoustic input (Ayala *et al*., 2016; Wetekam *et al*., 2025), such as those occurring during vocalizations or when new acoustic objects enter the surrounding environment. Since T-stellate neurons synapse onto inhibitory NPY neurons and excitatory VIP neurons, both of which make local axon collaterals (Goyer *et al*., 2019; Beebe *et al*., 2022; Silveira *et al*., 2024), T-stellate input likely recruits both inhibitory and excitatory local circuits in the IC. Interactions with local circuits may further enhance the detection of changes in sensory input. Currently, these are difficult predictions to test since the organization of IC local circuits remains largely unknown (Drotos & Roberts, 2024), but as IC studies advance, it will be exciting to test whether short-term plasticity and IC circuits generate filters for detecting sensory changes like those found in fish torus semicircularis (Fortune & Rose, 2000) and mouse cerebellum (Holla *et al*., 2026).

## Supporting information

Supplemental Statistics Tables

## Data Availability

Data will be made available upon reasonable request.

## Author contributions

YNH and MTR conceived and designed research. YNH and BMA performed experiments. YNH analyzed data. YNH and MTR interpreted results of experiments, prepared figures, drafted manuscript, and edited and revised manuscript. All authors approved the final version of the manuscript.

## Acknowledgements

We thank Pierre Apostolides, Marina Silveira, and Audrey Drotos for helpful discussions and advice. This work was supported by National Institutes of Health Grants F31 DC021618 (YNH), R25 NS107159 (BMA, PI: RK Duncan), and R01 DC018284 (MTR). Keywords – T-stellate neurons, cochlear nucleus, auditory system, short-term plasticity, inferior colliculus, electrophysiology, neural circuits

