## Supplemental Statistics Tables for "Optogenetic circuit mapping reveals connectivity and synaptic physiology of T-stellate projections from the cochlear nucleus to the auditory midbrain"

**Supplemental Table 1: oEPSC amplitude linear mixed model output**. LMM pairwise comparisons of each oEPSC amplitude compared to the first at all stimulation frequencies across both neuron types.

| 20 Hz | | | | | | | | | | | | | | |
|---|---|---|---|---|---|---|---|---|---|---|---|---|---|---|
| | NPY | | | | | | | VIP | | | | | | |
| Peak | Estimate | Std Error | df | t-value | p.value | CI Low | CI High | Estimate | Std Error | df | t-value | p.value | CI Low | CI High |
| **Peak 1** | 1.00 | 0.06 | 13.49 | 17.22 | p < 0.001 | 0.89 | 1.11 | 1.00 | 0.05 | 16.98 | 20.44 | p < 0.001 | 0.90 | 1.10 |
| **Peak 2** | -0.11 | 0.05 | 63.00 | -2.40 | 0.019 | -0.20 | -0.03 | -0.22 | 0.04 | 72.00 | -5.25 | p < 0.001 | -0.29 | -0.14 |
| **Peak 3** | -0.22 | 0.05 | 63.00 | -4.83 | p < 0.001 | -0.31 | -0.14 | -0.26 | 0.04 | 72.00 | -6.24 | p < 0.001 | -0.34 | -0.18 |
| **Peak 4** | -0.26 | 0.05 | 63.00 | -5.62 | p < 0.001 | -0.35 | -0.17 | -0.28 | 0.04 | 72.00 | -6.86 | p < 0.001 | -0.36 | -0.21 |
| **Peak 5** | -0.29 | 0.05 | 63.00 | -6.30 | p < 0.001 | -0.38 | -0.21 | -0.32 | 0.04 | 72.00 | -7.69 | p < 0.001 | -0.40 | -0.24 |
| **Peak 6** | -0.34 | 0.05 | 63.00 | -7.28 | p < 0.001 | -0.42 | -0.25 | -0.31 | 0.04 | 72.00 | -7.46 | p < 0.001 | -0.39 | -0.23 |
| **Peak 7** | -0.35 | 0.05 | 63.00 | -7.46 | p < 0.001 | -0.43 | -0.26 | -0.34 | 0.04 | 72.00 | -8.14 | p < 0.001 | -0.41 | -0.26 |
| **Peak 8** | -0.32 | 0.05 | 63.00 | -6.91 | p < 0.001 | -0.41 | -0.23 | -0.37 | 0.04 | 72.00 | -9.03 | p < 0.001 | -0.45 | -0.30 |
| **Peak 9** | -0.42 | 0.05 | 63.00 | -8.98 | p < 0.001 | -0.50 | -0.33 | -0.40 | 0.04 | 72.00 | -9.75 | p < 0.001 | -0.48 | -0.33 |
| **Peak 10** | -0.38 | 0.05 | 63.00 | -8.21 | p < 0.001 | -0.47 | -0.29 | -0.35 | 0.04 | 72.00 | -8.55 | p < 0.001 | -0.43 | -0.28 |
| 50 Hz | | | | | | | | | | | | | | |
| | NPY | | | | | | | VIP | | | | | | |
| Peak | Estimate | Std Error | df | t-value | p.value | CI Low | CI High | Estimate | Std Error | df | t-value | p.value | CI Low | CI High |
| **Peak 1** | 1.00 | 0.05 | 19.54 | 21.49 | p < 0.001 | 0.91 | 1.09 | 1.00 | 0.06 | 13.91 | 16.77 | p < 0.001 | 0.88 | 1.12 |
| **Peak 2** | -0.25 | 0.04 | 63.00 | -5.67 | p < 0.001 | -0.34 | -0.17 | -0.25 | 0.04 | 72.00 | -5.77 | p < 0.001 | -0.34 | -0.17 |
| **Peak 3** | -0.38 | 0.04 | 63.00 | -8.52 | p < 0.001 | -0.47 | -0.30 | -0.31 | 0.04 | 72.00 | -6.92 | p < 0.001 | -0.39 | -0.22 |
| **Peak 4** | -0.41 | 0.04 | 63.00 | -9.05 | p < 0.001 | -0.49 | -0.32 | -0.39 | 0.04 | 72.00 | -8.76 | p < 0.001 | -0.47 | -0.30 |
| **Peak 5** | -0.44 | 0.04 | 63.00 | -9.77 | p < 0.001 | -0.52 | -0.35 | -0.39 | 0.04 | 72.00 | -8.82 | p < 0.001 | -0.47 | -0.31 |
| **Peak 6** | -0.47 | 0.04 | 63.00 | -10.44 | p < 0.001 | -0.55 | -0.38 | -0.44 | 0.04 | 72.00 | -10.06 | p < 0.001 | -0.53 | -0.36 |
| **Peak 7** | -0.46 | 0.04 | 63.00 | -10.21 | p < 0.001 | -0.54 | -0.37 | -0.44 | 0.04 | 72.00 | -9.95 | p < 0.001 | -0.52 | -0.36 |
| **Peak 8** | -0.48 | 0.04 | 63.00 | -10.73 | p < 0.001 | -0.56 | -0.40 | -0.49 | 0.04 | 72.00 | -11.09 | p < 0.001 | -0.57 | -0.41 |
| **Peak 9** | -0.51 | 0.04 | 63.00 | -11.39 | p < 0.001 | -0.59 | -0.43 | -0.45 | 0.04 | 72.00 | -10.20 | p < 0.001 | -0.53 | -0.37 |
| **Peak 10** | -0.50 | 0.04 | 63.00 | -11.06 | p < 0.001 | -0.58 | -0.41 | -0.45 | 0.04 | 72.00 | -10.29 | p < 0.001 | -0.54 | -0.37 |
| 70 Hz | | | | | | | | | | | | | | |
| | NPY | | | | | | | VIP | | | | | | |
| Peak | Estimate | Std Error | df | t-value | p.value | CI Low | CI High | Estimate | Std Error | df | t-value | p.value | CI Low | CI High |
| **Peak 1** | 1.00 | 0.09 | 6.37 | 11.13 | p < 0.001 | 0.82 | 1.18 | 1.00 | 0.08 | 8.72 | 12.41 | p < 0.001 | 0.84 | 1.16 |
| **Peak 2** | -0.22 | 0.06 | 36.00 | -3.50 | 0.001 | -0.33 | -0.11 | -0.25 | 0.05 | 54.00 | -4.94 | p < 0.001 | -0.34 | -0.15 |
| **Peak 3** | -0.38 | 0.06 | 36.00 | -6.11 | p < 0.001 | -0.49 | -0.27 | -0.38 | 0.05 | 54.00 | -7.59 | p < 0.001 | -0.47 | -0.29 |
| **Peak 4** | -0.42 | 0.06 | 36.00 | -6.81 | p < 0.001 | -0.53 | -0.31 | -0.45 | 0.05 | 54.00 | -9.09 | p < 0.001 | -0.55 | -0.36 |
| **Peak 5** | -0.42 | 0.06 | 36.00 | -6.85 | p < 0.001 | -0.53 | -0.31 | -0.48 | 0.05 | 54.00 | -9.63 | p < 0.001 | -0.57 | -0.39 |
| **Peak 6** | -0.47 | 0.06 | 36.00 | -7.66 | p < 0.001 | -0.58 | -0.36 | -0.48 | 0.05 | 54.00 | -9.68 | p < 0.001 | -0.57 | -0.39 |
| **Peak 7** | -0.52 | 0.06 | 36.00 | -8.45 | p < 0.001 | -0.63 | -0.41 | -0.50 | 0.05 | 54.00 | -9.94 | p < 0.001 | -0.59 | -0.40 |
| **Peak 8** | -0.53 | 0.06 | 36.00 | -8.59 | p < 0.001 | -0.64 | -0.42 | -0.52 | 0.05 | 54.00 | -10.45 | p < 0.001 | -0.61 | -0.43 |
| **Peak 9** | -0.54 | 0.06 | 36.00 | -8.74 | p < 0.001 | -0.65 | -0.43 | -0.55 | 0.05 | 54.00 | -11.00 | p < 0.001 | -0.64 | -0.46 |
| **Peak 10** | -0.54 | 0.06 | 36.00 | -8.80 | p < 0.001 | -0.65 | -0.43 | -0.54 | 0.05 | 54.00 | -10.83 | p < 0.001 | -0.63 | -0.45 |

**Supplemental Table 2: AAV9 location of activation pairwise comparisons**. LMM pairwise comparisons of each oEPSC latency compared to the first at each location of activation for AAV9 viral serotype control.

| AAV9 on Bouton | | | | | | | |
|---|---|---|---|---|---|---|---|
| Peak | Estimate | Std Error | df | t-value | p.value | CI Low | CI High |
| **Peak 1** | 1.00 | 0.08 | 16.51 | 13.32 | p < 0.001 | 0.85 | 1.15 |
| **Peak 2** | -0.14 | 0.07 | 63.00 | -2.09 | 0.041 | -0.27 | -0.02 |
| **Peak 3** | -0.27 | 0.07 | 63.00 | -4.03 | p < 0.001 | -0.40 | -0.15 |
| **Peak 4** | -0.22 | 0.07 | 63.00 | -3.34 | 0.001 | -0.35 | -0.10 |
| **Peak 5** | -0.28 | 0.07 | 63.00 | -4.17 | p < 0.001 | -0.40 | -0.16 |
| **Peak 6** | -0.29 | 0.07 | 63.00 | -4.33 | p < 0.001 | -0.42 | -0.17 |
| **Peak 7** | -0.34 | 0.07 | 63.00 | -5.12 | p < 0.001 | -0.47 | -0.22 |
| **Peak 8** | -0.27 | 0.07 | 63.00 | -3.97 | p < 0.001 | -0.39 | -0.14 |
| **Peak 9** | -0.30 | 0.07 | 63.00 | -4.48 | p < 0.001 | -0.43 | -0.18 |
| **Peak 10** | -0.35 | 0.07 | 63.00 | -5.28 | p < 0.001 | -0.48 | -0.23 |
| AAV9 off Bouton | | | | | | | |
| Peak | Estimate | Std Error | df | t-value | p.value | CI Low | CI High |
| **Peak 1** | 1.00 | 0.06 | 21.40 | 15.90 | p < 0.001 | 0.88 | 1.12 |
| **Peak 2** | -0.12 | 0.06 | 63.00 | -1.95 | 0.056 | -0.24 | -0.01 |
| **Peak 3** | -0.20 | 0.06 | 63.00 | -3.15 | 0.002 | -0.31 | -0.08 |
| **Peak 4** | -0.27 | 0.06 | 63.00 | -4.33 | p < 0.001 | -0.39 | -0.16 |
| **Peak 5** | -0.28 | 0.06 | 63.00 | -4.45 | p < 0.001 | -0.40 | -0.16 |
| **Peak 6** | -0.27 | 0.06 | 63.00 | -4.35 | p < 0.001 | -0.39 | -0.16 |
| **Peak 7** | -0.26 | 0.06 | 63.00 | -4.07 | p < 0.001 | -0.37 | -0.14 |
| **Peak 8** | -0.19 | 0.06 | 63.00 | -3.03 | 0.004 | -0.31 | -0.07 |
| **Peak 9** | -0.21 | 0.06 | 63.00 | -3.27 | 0.002 | -0.32 | -0.09 |
| **Peak 10** | -0.29 | 0.06 | 63.00 | -4.60 | p < 0.001 | -0.40 | -0.17 |

**Supplemental Table 3: oEPSC failure rate linear mixed model output**. LMM pairwise comparisons of each oEPSC failure rate compared to the first at all stimulation frequencies across both neuron types.

| 20 Hz | | | | | | | | | | | | | | |
|---|---|---|---|---|---|---|---|---|---|---|---|---|---|---|
| | NPY | | | | | | | VIP | | | | | | |
| Peak | Estimate | Std Error | df | t-value | p.value | CI Low | CI High | Estimate | Std Error | df | t-value | p.value | CI Low | CI High |
| **Peak 1** | 0.03 | 0.05 | 20.11 | 0.58 | 0.567 | -0.07 | 0.13 | 0.07 | 0.07 | 12.39 | 1.095 | 0.294 | -0.06 | 0.208 |
| **Peak 2** | 0.09 | 0.05 | 63.00 | 1.67 | 0.100 | -0.01 | 0.18 | 0.10 | 0.04 | 72.00 | 2.267 | 0.026 | 0.02 | 0.185 |
| **Peak 3** | 0.12 | 0.05 | 63.00 | 2.39 | 0.020 | 0.03 | 0.22 | 0.14 | 0.04 | 72.00 | 3.168 | 0.002 | 0.06 | 0.225 |
| **Peak 4** | 0.20 | 0.05 | 63.00 | 3.82 | $p < 0.001$ | 0.10 | 0.30 | 0.17 | 0.04 | 72.00 | 3.758 | $p < 0.001$ | 0.08 | 0.252 |
| **Peak 5** | 0.17 | 0.05 | 63.00 | 3.22 | 0.002 | 0.07 | 0.27 | 0.20 | 0.04 | 72.00 | 4.410 | $p < 0.001$ | 0.11 | 0.281 |
| **Peak 6** | 0.16 | 0.05 | 63.00 | 2.98 | 0.004 | 0.06 | 0.25 | 0.17 | 0.04 | 72.00 | 3.851 | $p < 0.001$ | 0.09 | 0.256 |
| **Peak 7** | 0.22 | 0.05 | 63.00 | 4.30 | $p < 0.001$ | 0.13 | 0.32 | 0.20 | 0.04 | 72.00 | 4.503 | $p < 0.001$ | 0.12 | 0.285 |
| **Peak 8** | 0.26 | 0.05 | 63.00 | 4.89 | $p < 0.001$ | 0.16 | 0.35 | 0.25 | 0.04 | 72.00 | 5.653 | $p < 0.001$ | 0.17 | 0.336 |
| **Peak 9** | 0.19 | 0.05 | 63.00 | 3.58 | $p < 0.001$ | 0.09 | 0.28 | 0.27 | 0.04 | 72.00 | 6.149 | $p < 0.001$ | 0.19 | 0.359 |
| **Peak 10** | 0.24 | 0.05 | 63.00 | 4.66 | $p < 0.001$ | 0.15 | 0.34 | 0.30 | 0.04 | 72.00 | 6.615 | $p < 0.001$ | 0.21 | 0.379 |

| 50 Hz | | | | | | | | | | | | | | |
|---|---|---|---|---|---|---|---|---|---|---|---|---|---|---|
| | NPY | | | | | | | VIP | | | | | | |
| Peak | Estimate | Std Error | df | t-value | p.value | CI Low | CI High | Estimate | Std Error | df | t-value | p.value | CI Low | CI High |
| **Peak 1** | 0.06 | 0.06 | 25.59 | 1.03 | 0.384 | -0.05 | 0.18 | 0.07 | 0.07 | 11.26 | 1.11 | 0.053 | -0.06 | 0.21 |
| **Peak 2** | 0.06 | 0.06 | 63.00 | 0.88 | 0.029 | -0.06 | 0.18 | 0.08 | 0.04 | 72.00 | 1.97 | $p < 0.001$ | 0.00 | 0.15 |
| **Peak 3** | 0.14 | 0.06 | 63.00 | 2.24 | 0.210 | 0.02 | 0.26 | 0.16 | 0.04 | 72.00 | 3.99 | $p < 0.001$ | 0.08 | 0.23 |
| **Peak 4** | 0.08 | 0.06 | 63.00 | 1.27 | 0.045 | -0.04 | 0.20 | 0.14 | 0.04 | 72.00 | 3.57 | $p < 0.001$ | 0.07 | 0.21 |
| **Peak 5** | 0.13 | 0.06 | 63.00 | 2.04 | 0.005 | 0.01 | 0.25 | 0.24 | 0.04 | 72.00 | 6.16 | $p < 0.001$ | 0.17 | 0.31 |
| **Peak 6** | 0.19 | 0.06 | 63.00 | 2.92 | 0.002 | 0.07 | 0.31 | 0.22 | 0.04 | 72.00 | 5.59 | $p < 0.001$ | 0.15 | 0.29 |
| **Peak 7** | 0.21 | 0.06 | 63.00 | 3.21 | $p < 0.001$ | 0.09 | 0.33 | 0.30 | 0.04 | 72.00 | 7.58 | $p < 0.001$ | 0.22 | 0.37 |
| **Peak 8** | 0.24 | 0.06 | 63.00 | 3.70 | 0.002 | 0.12 | 0.36 | 0.27 | 0.04 | 72.00 | 6.88 | $p < 0.001$ | 0.20 | 0.34 |
| **Peak 9** | 0.21 | 0.06 | 63.00 | 3.31 | $p < 0.001$ | 0.09 | 0.33 | 0.30 | 0.04 | 72.00 | 7.59 | $p < 0.001$ | 0.22 | 0.37 |
| **Peak 10** | 0.25 | 0.06 | 63.00 | 3.89 | $p < 0.001$ | 0.13 | 0.37 | 0.32 | 0.04 | 72.00 | 8.15 | $p < 0.001$ | 0.25 | 0.39 |

| 70 Hz | | | | | | | | | | | | | | |
|---|---|---|---|---|---|---|---|---|---|---|---|---|---|---|
| | NPY | | | | | | | VIP | | | | | | |
| Peak | Estimate | Std Error | df | t-value | p.value | CI Low | CI High | Estimate | Std Error | df | t-value | p.value | CI Low | CI High |
| **Peak 1** | 0.05 | 0.06 | 16.40 | 0.88 | $p < 0.001$ | -0.06 | 0.16 | 0.06 | 0.05 | 17.11 | 1.08 | 0.295 | -0.04 | 0.16 |
| **Peak 2** | 0.07 | 0.06 | 36.00 | 1.12 | $p < 0.001$ | -0.04 | 0.18 | 0.02 | 0.05 | 54.00 | 0.42 | 0.679 | -0.07 | 0.12 |
| **Peak 3** | 0.12 | 0.06 | 36.00 | 1.93 | $p < 0.001$ | 0.01 | 0.23 | 0.10 | 0.05 | 54.00 | 1.94 | 0.057 | 0.01 | 0.19 |
| **Peak 4** | 0.15 | 0.06 | 36.00 | 2.41 | $p < 0.001$ | 0.04 | 0.26 | 0.10 | 0.05 | 54.00 | 1.94 | 0.057 | 0.01 | 0.19 |
| **Peak 5** | 0.24 | 0.06 | 36.00 | 3.85 | $p < 0.001$ | 0.13 | 0.35 | 0.11 | 0.05 | 54.00 | 2.08 | 0.042 | 0.01 | 0.20 |
| **Peak 6** | 0.15 | 0.06 | 36.00 | 2.41 | $p < 0.001$ | 0.04 | 0.26 | 0.14 | 0.05 | 54.00 | 2.78 | 0.008 | 0.05 | 0.24 |
| **Peak 7** | 0.17 | 0.06 | 36.00 | 2.73 | $p < 0.001$ | 0.06 | 0.28 | 0.21 | 0.05 | 54.00 | 4.17 | $p < 0.001$ | 0.12 | 0.31 |
| **Peak 8** | 0.20 | 0.06 | 36.00 | 3.21 | $p < 0.001$ | 0.09 | 0.31 | 0.24 | 0.05 | 54.00 | 4.58 | $p < 0.001$ | 0.14 | 0.33 |
| **Peak 9** | 0.22 | 0.06 | 36.00 | 3.53 | $p < 0.001$ | 0.11 | 0.33 | 0.19 | 0.05 | 54.00 | 3.75 | $p < 0.001$ | 0.10 | 0.29 |
| **Peak 10** | 0.23 | 0.06 | 36.00 | 3.69 | $p < 0.001$ | 0.12 | 0.34 | 0.24 | 0.05 | 54.00 | 4.72 | $p < 0.001$ | 0.15 | 0.34 |

**Supplemental Table 4: oEPSC latency linear mixed model output**. LMM pairwise comparisons of each oEPSC latency compared to the first at all stimulation frequencies across both neuron types.

| 20 Hz | | | | | | | | | | | | | | |
|---|---|---|---|---|---|---|---|---|---|---|---|---|---|---|
| | NPY | | | | | | | VIP | | | | | | |
| Peak | Estimate | Std Error | df | t-value | p.value | CI Low | CI High | Estimate | Std Error | df | t-value | p.value | CI Low | CI High |
| **Peak 1** | 1.91 | 0.18 | 8.38 | 10.89 | p < 0.001 | 1.55 | 2.26 | 1.97 | 0.17 | 9.52 | 11.58 | p < 0.001 | 1.62 | 2.31 |
| **Peak 2** | 0.15 | 0.08 | 63.00 | 1.99 | 0.051 | 0.01 | 0.30 | 0.20 | 0.07 | 72.00 | 2.76 | 0.007 | 0.06 | 0.34 |
| **Peak 3** | 0.16 | 0.08 | 63.00 | 2.06 | 0.043 | 0.02 | 0.30 | 0.33 | 0.07 | 72.00 | 4.49 | p < 0.001 | 0.19 | 0.47 |
| **Peak 4** | 0.21 | 0.08 | 63.00 | 2.78 | 0.007 | 0.07 | 0.36 | 0.32 | 0.07 | 72.00 | 4.33 | p < 0.001 | 0.18 | 0.45 |
| **Peak 5** | 0.15 | 0.08 | 63.00 | 1.98 | 0.052 | 0.01 | 0.29 | 0.39 | 0.07 | 72.00 | 5.32 | p < 0.001 | 0.25 | 0.53 |
| **Peak 6** | 0.31 | 0.08 | 63.00 | 3.98 | p < 0.001 | 0.16 | 0.45 | 0.39 | 0.07 | 72.00 | 5.29 | p < 0.001 | 0.25 | 0.52 |
| **Peak 7** | 0.30 | 0.08 | 63.00 | 3.86 | p < 0.001 | 0.15 | 0.44 | 0.40 | 0.07 | 72.00 | 5.41 | p < 0.001 | 0.26 | 0.53 |
| **Peak 8** | 0.28 | 0.08 | 63.00 | 3.65 | p < 0.001 | 0.14 | 0.42 | 0.49 | 0.07 | 72.00 | 6.68 | p < 0.001 | 0.35 | 0.63 |
| **Peak 9** | 0.49 | 0.08 | 63.00 | 6.43 | p < 0.001 | 0.35 | 0.64 | 0.52 | 0.07 | 72.00 | 7.13 | p < 0.001 | 0.39 | 0.66 |
| **Peak 10** | 0.38 | 0.08 | 63.00 | 4.89 | p < 0.001 | 0.23 | 0.52 | 0.39 | 0.07 | 72.00 | 5.33 | p < 0.001 | 0.25 | 0.53 |
| 50 Hz | | | | | | | | | | | | | | |
| | NPY | | | | | | | VIP | | | | | | |
| Peak | Estimate | Std Error | df | t-value | p.value | CI Low | CI High | Estimate | Std Error | df | t-value | p.value | CI Low | CI High |
| **Peak 1** | 1.86 | 0.18 | 9.17 | 10.46 | p < 0.001 | 1.50 | 2.22 | 1.83 | 0.18 | 9.68 | 10.17 | p < 0.001 | 1.47 | 2.20 |
| **Peak 2** | 0.12 | 0.09 | 63.00 | 1.31 | 0.196 | -0.05 | 0.30 | 0.33 | 0.08 | 72.00 | 4.05 | p < 0.001 | 0.18 | 0.48 |
| **Peak 3** | 0.21 | 0.09 | 63.00 | 2.27 | 0.027 | 0.04 | 0.39 | 0.44 | 0.08 | 72.00 | 5.40 | p < 0.001 | 0.29 | 0.59 |
| **Peak 4** | 0.34 | 0.09 | 63.00 | 3.61 | p < 0.001 | 0.17 | 0.52 | 0.54 | 0.08 | 72.00 | 6.60 | p < 0.001 | 0.38 | 0.69 |
| **Peak 5** | 0.41 | 0.09 | 63.00 | 4.33 | p < 0.001 | 0.23 | 0.58 | 0.43 | 0.08 | 72.00 | 5.24 | p < 0.001 | 0.27 | 0.58 |
| **Peak 6** | 0.34 | 0.09 | 63.00 | 3.57 | p < 0.001 | 0.16 | 0.51 | 0.60 | 0.08 | 72.00 | 7.38 | p < 0.001 | 0.45 | 0.75 |
| **Peak 7** | 0.38 | 0.09 | 63.00 | 4.00 | p < 0.001 | 0.20 | 0.55 | 0.54 | 0.08 | 72.00 | 6.63 | p < 0.001 | 0.39 | 0.69 |
| **Peak 8** | 0.41 | 0.09 | 63.00 | 4.39 | p < 0.001 | 0.24 | 0.59 | 0.60 | 0.08 | 72.00 | 7.36 | p < 0.001 | 0.45 | 0.75 |
| **Peak 9** | 0.44 | 0.09 | 63.00 | 4.63 | p < 0.001 | 0.26 | 0.61 | 0.67 | 0.08 | 72.00 | 8.24 | p < 0.001 | 0.52 | 0.82 |
| **Peak 10** | 0.48 | 0.09 | 63.00 | 5.10 | p < 0.001 | 0.31 | 0.66 | 0.66 | 0.08 | 72.00 | 8.11 | p < 0.001 | 0.51 | 0.81 |
| 70 Hz | | | | | | | | | | | | | | |
| | NPY | | | | | | | VIP | | | | | | |
| Peak | Estimate | Std Error | df | t-value | p.value | CI Low | CI High | Estimate | Std Error | df | t-value | p.value | CI Low | CI High |
| **Peak 1** | 1.92 | 0.24 | 5.82 | 7.94 | p < 0.001 | 1.42 | 2.42 | 2.20 | 0.26 | 7.21 | 8.30 | p < 0.001 | 1.65 | 2.74 |
| **Peak 2** | 0.07 | 0.15 | 36.00 | 0.47 | 0.642 | -0.20 | 0.34 | 0.20 | 0.12 | 54.00 | 1.71 | 0.094 | -0.02 | 0.42 |
| **Peak 3** | 0.17 | 0.15 | 36.00 | 1.14 | 0.262 | -0.10 | 0.44 | 0.30 | 0.12 | 54.00 | 2.57 | 0.013 | 0.08 | 0.52 |
| **Peak 4** | 0.28 | 0.15 | 36.00 | 1.90 | 0.066 | 0.02 | 0.55 | 0.35 | 0.12 | 54.00 | 3.01 | 0.004 | 0.14 | 0.57 |
| **Peak 5** | 0.28 | 0.15 | 36.00 | 1.86 | 0.071 | 0.01 | 0.55 | 0.35 | 0.12 | 54.00 | 2.98 | 0.004 | 0.13 | 0.57 |
| **Peak 6** | 0.44 | 0.15 | 36.00 | 2.95 | 0.006 | 0.17 | 0.71 | 0.44 | 0.12 | 54.00 | 3.77 | p < 0.001 | 0.23 | 0.66 |
| **Peak 7** | 0.35 | 0.15 | 36.00 | 2.31 | 0.027 | 0.08 | 0.62 | 0.58 | 0.12 | 54.00 | 4.92 | p < 0.001 | 0.36 | 0.79 |
| **Peak 8** | 0.30 | 0.15 | 36.00 | 2.02 | 0.051 | 0.03 | 0.57 | 0.54 | 0.12 | 54.00 | 4.62 | p < 0.001 | 0.33 | 0.76 |
| **Peak 9** | 0.38 | 0.15 | 36.00 | 2.56 | 0.015 | 0.11 | 0.65 | 0.58 | 0.12 | 54.00 | 4.95 | p < 0.001 | 0.36 | 0.80 |
| **Peak 10** | 0.44 | 0.15 | 36.00 | 2.95 | 0.005 | 0.17 | 0.71 | 0.55 | 0.12 | 54.00 | 4.71 | p < 0.001 | 0.34 | 0.77 |

**Supplemental Table 5: oEPSC jitter linear mixed model output**. LMM pairwise comparisons of each oEPSC jitter compared to the first at all stimulation frequencies across both neuron types.

| 20 Hz | | | | | | | | | | | | | | |
|---|---|---|---|---|---|---|---|---|---|---|---|---|---|---|
| | NPY | | | | | | | VIP | | | | | | |
| Peak | Estimate | Std Error | df | t-value | p.value | CI Low | CI High | Estimate | Std Error | df | t-value | p.value | CI Low | CI High |
| **Peak 1** | 0.44 | 0.07 | 11.87 | 6.11 | p < 0.001 | 0.30 | 0.58 | 0.28 | 0.06 | 40.95 | 4.88 | p < 0.001 | 0.17 | 0.39 |
| **Peak 2** | -0.02 | 0.05 | 63.00 | -0.33 | 0.740 | -0.11 | 0.08 | 0.10 | 0.07 | 72.00 | 1.46 | 0.148 | -0.03 | 0.22 |
| **Peak 3** | -0.02 | 0.05 | 63.00 | -0.40 | 0.690 | -0.12 | 0.08 | 0.16 | 0.07 | 72.00 | 2.42 | 0.018 | 0.04 | 0.29 |
| **Peak 4** | -0.02 | 0.05 | 63.00 | -0.42 | 0.678 | -0.12 | 0.07 | 0.15 | 0.07 | 72.00 | 2.26 | 0.027 | 0.03 | 0.27 |
| **Peak 5** | 0.01 | 0.05 | 63.00 | 0.28 | 0.782 | -0.08 | 0.11 | 0.16 | 0.07 | 72.00 | 2.38 | 0.020 | 0.03 | 0.28 |
| **Peak 6** | 0.03 | 0.05 | 63.00 | 0.60 | 0.549 | -0.07 | 0.13 | 0.19 | 0.07 | 72.00 | 2.85 | 0.006 | 0.06 | 0.31 |
| **Peak 7** | -0.02 | 0.05 | 63.00 | -0.36 | 0.718 | -0.12 | 0.08 | 0.23 | 0.07 | 72.00 | 3.45 | p < 0.001 | 0.10 | 0.35 |
| **Peak 8** | 0.07 | 0.05 | 63.00 | 1.35 | 0.182 | -0.03 | 0.17 | 0.20 | 0.07 | 72.00 | 3.06 | 0.003 | 0.08 | 0.33 |
| **Peak 9** | 0.10 | 0.05 | 63.00 | 1.98 | 0.052 | 0.01 | 0.20 | 0.21 | 0.07 | 72.00 | 3.20 | 0.002 | 0.09 | 0.34 |
| **Peak 10** | 0.09 | 0.05 | 63.00 | 1.66 | 0.101 | -0.01 | 0.18 | 0.11 | 0.07 | 72.00 | 1.72 | 0.090 | -0.01 | 0.24 |
| 50 Hz | | | | | | | | | | | | | | |
| | NPY | | | | | | | VIP | | | | | | |
| Peak | Estimate | Std Error | df | t-value | p.value | CI Low | CI High | Estimate | Std Error | df | t-value | p.value | CI Low | CI High |
| **Peak 1** | 0.36 | 0.09 | 19.69 | 4.24 | p < 0.001 | 0.20 | 0.53 | 0.19 | 0.05 | 66.60 | 3.62 | p < 0.001 | 0.09 | 0.29 |
| **Peak 2** | 0.00 | 0.08 | 63.00 | -0.03 | 0.979 | -0.16 | 0.15 | 0.15 | 0.07 | 72.00 | 2.12 | 0.037 | 0.02 | 0.28 |
| **Peak 3** | 0.05 | 0.08 | 63.00 | 0.60 | 0.552 | -0.10 | 0.20 | 0.17 | 0.07 | 72.00 | 2.40 | 0.019 | 0.04 | 0.30 |
| **Peak 4** | 0.09 | 0.08 | 63.00 | 1.14 | 0.259 | -0.06 | 0.25 | 0.26 | 0.07 | 72.00 | 3.81 | p < 0.001 | 0.13 | 0.39 |
| **Peak 5** | 0.13 | 0.08 | 63.00 | 1.62 | 0.111 | -0.02 | 0.29 | 0.13 | 0.07 | 72.00 | 1.82 | 0.073 | 0.00 | 0.26 |
| **Peak 6** | 0.19 | 0.08 | 63.00 | 2.33 | 0.023 | 0.04 | 0.35 | 0.19 | 0.07 | 72.00 | 2.68 | 0.009 | 0.06 | 0.31 |
| **Peak 7** | 0.09 | 0.08 | 63.00 | 1.11 | 0.273 | -0.06 | 0.24 | 0.18 | 0.07 | 72.00 | 2.59 | 0.012 | 0.05 | 0.31 |
| **Peak 8** | 0.10 | 0.08 | 63.00 | 1.20 | 0.234 | -0.05 | 0.25 | 0.25 | 0.07 | 72.00 | 3.58 | p < 0.001 | 0.12 | 0.38 |
| **Peak 9** | 0.07 | 0.08 | 63.00 | 0.83 | 0.412 | -0.09 | 0.22 | 0.21 | 0.07 | 72.00 | 3.09 | 0.003 | 0.08 | 0.34 |
| **Peak 10** | 0.19 | 0.08 | 63.00 | 2.27 | 0.027 | 0.03 | 0.34 | 0.17 | 0.07 | 72.00 | 2.50 | 0.015 | 0.04 | 0.30 |
| 70 Hz | | | | | | | | | | | | | | |
| | NPY | | | | | | | VIP | | | | | | |
| Peak | Estimate | Std Error | df | t-value | p.value | CI Low | CI High | Estimate | Std Error | df | t-value | p.value | CI Low | CI High |
| **Peak 1** | 0.23 | 0.07 | 11.78 | 3.27 | 0.007 | 0.10 | 0.37 | 0.34 | 0.07 | 54.84 | 5.20 | p < 0.001 | 0.22 | 0.47 |
| **Peak 2** | 0.04 | 0.07 | 36.00 | 0.55 | 0.588 | -0.09 | 0.16 | -0.04 | 0.09 | 54.00 | -0.44 | 0.664 | -0.20 | 0.12 |
| **Peak 3** | 0.06 | 0.07 | 36.00 | 0.89 | 0.381 | -0.06 | 0.19 | 0.07 | 0.09 | 54.00 | 0.84 | 0.403 | -0.09 | 0.24 |
| **Peak 4** | 0.09 | 0.07 | 36.00 | 1.34 | 0.188 | -0.03 | 0.22 | 0.00 | 0.09 | 54.00 | 0.00 | 0.998 | -0.16 | 0.16 |
| **Peak 5** | 0.15 | 0.07 | 36.00 | 2.06 | 0.046 | 0.02 | 0.27 | -0.09 | 0.09 | 54.00 | -1.04 | 0.304 | -0.26 | 0.07 |
| **Peak 6** | 0.11 | 0.07 | 36.00 | 1.63 | 0.113 | -0.01 | 0.24 | 0.11 | 0.09 | 54.00 | 1.26 | 0.215 | -0.05 | 0.27 |
| **Peak 7** | 0.11 | 0.07 | 36.00 | 1.51 | 0.140 | -0.02 | 0.23 | 0.15 | 0.09 | 54.00 | 1.64 | 0.106 | -0.02 | 0.31 |
| **Peak 8** | 0.02 | 0.07 | 36.00 | 0.21 | 0.832 | -0.11 | 0.14 | 0.26 | 0.09 | 54.00 | 2.97 | 0.004 | 0.10 | 0.43 |
| **Peak 9** | 0.08 | 0.07 | 36.00 | 1.15 | 0.257 | -0.05 | 0.21 | 0.10 | 0.09 | 54.00 | 1.08 | 0.286 | -0.07 | 0.26 |
| **Peak 10** | 0.13 | 0.07 | 36.00 | 1.83 | 0.076 | 0.00 | 0.26 | -0.01 | 0.09 | 54.00 | -0.11 | 0.911 | -0.17 | 0.15 |

**Supplemental Table 6: oEPSC frequency pairwise comparisons**. LMM pairwise comparisons of neuron type oEPSC amplitude, failure rate, latency, and jitter pairwise comparisons of all frequencies of activation across both neuron types.

| Amplitude | | | | | | | | | | | | | | |
|---|---|---|---|---|---|---|---|---|---|---|---|---|---|---|
| | NPY | | | | | | | VIP | | | | | | |
| Frequency | Estimate | Std Error | df | t-ratio | p.value | CI Low | CI High | Estimate | Std Error | df | t-value | p.value | CI Low | CI High |
| **20-50** | 0.12 | 0.02 | 191 | 7.16 | $p < 0.001$ | 0.08 | 0.16 | 0.08 | 0.01 | 230 | 5.11 | $p < 0.001$ | 0.04 | 0.11 |
| **20-70** | 0.16 | 0.02 | 193 | 7.82 | $p < 0.001$ | 0.11 | 0.21 | 0.11 | 0.02 | 231 | 6.61 | $p < 0.001$ | 0.07 | 0.15 |
| **50-70** | 0.04 | 0.02 | 193 | 1.88 | 0.149 | -0.01 | 0.09 | 0.03 | 0.02 | 231 | 1.97 | 0.123 | -0.01 | 0.07 |

| Failure Rate | | | | | | | | | | | | | | |
|---|---|---|---|---|---|---|---|---|---|---|---|---|---|---|
| | NPY | | | | | | | VIP | | | | | | |
| Frequency | Estimate | Std Error | df | t-value | p.value | CI Low | CI High | Estimate | Std Error | df | t-value | p.value | CI Low | CI High |
| **20-50** | -0.02 | 0.02 | 191 | -0.96 | 0.602 | -0.06 | 0.02 | -0.02 | 0.01 | 230 | -1.34 | 0.376 | -0.05 | 0.02 |
| **20-70** | 0.01 | 0.02 | 193 | 0.56 | 0.841 | -0.04 | 0.06 | 0.01 | 0.02 | 230 | 0.37 | 0.928 | -0.03 | 0.04 |
| **50-70** | 0.03 | 0.02 | 193 | 1.36 | 0.364 | -0.02 | 0.08 | 0.03 | 0.02 | 230 | 1.58 | 0.255 | -0.01 | 0.06 |

| Latency | | | | | | | | | | | | | | |
|---|---|---|---|---|---|---|---|---|---|---|---|---|---|---|
| | NPY | | | | | | | VIP | | | | | | |
| Frequency | Estimate | Std Error | df | t-value | p.value | CI Low | CI High | Estimate | Std Error | df | t-value | p.value | CI Low | CI High |
| **20-50** | -0.02 | 0.03 | 191 | -0.69 | 0.772 | -0.09 | 0.05 | 0.00 | 0.04 | 230 | -0.10 | 0.994 | -0.10 | 0.09 |
| **20-70** | 0.06 | 0.04 | 191 | 1.66 | 0.222 | -0.03 | 0.15 | -0.24 | 0.04 | 230 | -5.55 | $p < 0.001$ | -0.35 | -0.14 |
| **50-70** | 0.08 | 0.04 | 191 | 2.23 | 0.068 | 0.00 | 0.17 | -0.24 | 0.04 | 230 | -5.46 | $p < 0.001$ | -0.34 | -0.14 |

| Jitter | | | | | | | | | | | | | | |
|---|---|---|---|---|---|---|---|---|---|---|---|---|---|---|
| | NPY | | | | | | | VIP | | | | | | |
| Frequency | Estimate | Std Error | df | t-value | p.value | CI Low | CI High | Estimate | Std Error | df | t-value | p.value | CI Low | CI High |
| **20-50** | 0.01 | 0.02 | 191 | 0.32 | 0.944 | -0.05 | 0.06 | 0.07 | 0.02 | 230 | 2.94 | 0.010 | 0.01 | 0.12 |
| **20-70** | 0.11 | 0.03 | 192 | 3.98 | $p < 0.001$ | 0.04 | 0.17 | 0.04 | 0.03 | 235 | 1.48 | 0.301 | -0.02 | 0.10 |
| **50-70** | 0.10 | 0.03 | 192 | 3.71 | $p < 0.001$ | 0.04 | 0.16 | -0.03 | 0.03 | 235 | -1.20 | 0.456 | -0.09 | 0.03 |
